# Separable global and local beta burst dynamics in motor cortex of primates

**DOI:** 10.1101/2025.05.05.652217

**Authors:** Preeya Khanna, Behraz Farrokhi, Hoseok Choi, Sandon Griffin, Ian Heimbuch, Lisa Novik, Katherina Thiesen, John Morrison, Robert J Morecraft, Karunesh Ganguly

## Abstract

Sensorimotor beta band oscillations are known to modulate during normal movement control and abnormal beta modulation is linked to pathological bradykinesia. However, the functional differences between beta localized to one brain area versus beta synchronized across brain areas remains unclear. We monitored beta bursts in non-human primates, both neurotypical and stroke-impaired, during the performance of complex motor tasks. Across both groups of animals, we identified two distinct beta burst types: global bursts that tend to be synchronized across cortical and subcortical areas, and local bursts that tend to be confined to cortex. These two types exhibited distinct neural dynamics, with global bursts linked to reduced firing variability and overall slowed movements. In contrast, local bursts often occurred during the execution of complex behaviors, particularly during prehension. We found evidence for changes in the distribution of global and local bursts during recovery after stroke. In impaired animals early after stroke, global bursts predominated and were associated with reduced speed and impaired grasping. Notably, recovery of grasping was associated with a reduction in global bursts and an increase in local bursts, suggesting that local bursts may play an important role during prehension. Our findings reveal distinct roles of global and local beta bursts and indicate that the normalization of global and local burst timing tracks recovery of dexterity.

## Introduction

Beta band oscillations (∼13-30 Hz, hereafter “beta”) are ubiquitous throughout the sensorimotor primate brain. They are associated with normal movement control and somatosensory processing^1–3^ and are also implicated in the pathophysiology of Parkinson’s disease (PD) and stroke^4,5^. Beta in both the cortex and the subcortex (e.g., basal ganglia and thalamus) is well known to undergo a coordinated suppression during simple, isolated, fast movements^2–4,6^. In PD, abnormal increases in beta in both cortical and subcortical areas are linked to bradykinesia and akinesia. Causal experiments have further shown that increases in beta result in slowing of movements^7–10^. Moreover, after stroke, incomplete beta suppression is linked with impaired movement control^11^. Together, this implies that coordinated cortical-subcortical “global” beta is an anti-kinetic signal, reflecting neural firing patterns that are incompatible with movement generation^12^. However, there are a few studies that appear to contradict the notion that beta always occurs globally and that it exclusively reflects an anti-kinetic state. For example, during complex movements – sequential or grasping actions – cortical and subcortical beta can become uncoupled such that each brain area has its own “local” beta dynamics^13^, and cortical beta has been shown to persist throughout dynamic movements^14,15^. This raises the possibility, evident only through studying complex movements, that local and global beta oscillations may signal two distinct neural computations.

In this study, we sought to characterize the cortical and subcortical beta dynamics that occur during complex behaviors in both stroke-impaired and neurotypical non-human primates. Leveraging recent findings that beta-band activity is better represented by brief, transient ’bursts’ rather than sustained oscillations^13,16^, we discovered two distinct categories of beta bursts: global beta bursts that tend to occur synchronously across cortex and subcortex, and local beta bursts that tend to occur in a spatially limited area of cortex. Previous modeling and experimental evidence indicates that cortex can generate beta oscillations just with its own local circutry^17–19^, or in response to inputs from subcortical regions^20^, supporting this separation. By focusing on temporally precise beta burst events, we were able to study beta dynamics on a single trial level, enabling us to study the roles of global and local beta events during variable, complex behavior.

Our results from both stroke-impaired and healthy non-human primates performing complex movement tasks indicate that global and local bursts represent distinct neural computations: global bursts are associated with movement-inhibiting neural patterns while local bursts are associated with neural variability supporting the execution of complex behaviors. In post-stroke animals with impaired reach-to-grasp movement, mid-movement global beta bursts occurred frequently and were linked to slower movement speeds and impaired grasping. With the recovery of fast and reliable reach-to-grasp movements, we observed a reduction in mid-movement global beta bursts, but a striking increase in local cortical beta bursts. Analysis of cortical single-unit recordings revealed distinct roles: global bursts were associated with strong single-unit entrainment, and reduced firing rates and firing variability, whereas local bursts were spatially clustered in cortex and were associated with weak single-unit entrainment and preserved firing variability. Similar global and local patterns were observed in intact animals during complex sequential movement tasks. Altogether, these results highlight that global bursts are associated with constrained neural firing that is incompatible with fast movement, while local bursts are associated with greater neural variability and the ongoing execution of complex behaviors. These findings provide new insights into beta oscillations and their potential to understand recovery of motor control after stroke.

## Results

### Recovery of dexterity from an M1 stroke

To study cortical and subcortical signals during complex movement in impaired and recovered primates, we first trained rhesus macaques to perform a reach-to-grasp task (Fig. 1A, Fig. S1). Animals initiated trials by pressing a button, then reached out, grasped an object through a slot, and lifted and held it above a designated height for a designated duration. Grasp difficulty depended on slot shape, where the precision slot was the most difficult and often required multiple manipulation attempts and the power slot was the easiest (Fig. 1B). Animals were trained until performance plateaued. Next, animals underwent a stroke induction in which the one hemisphere’s primary motor cortex (M1) arm and hand area was targeted with surface vessel occlusion followed by subpial aspiration^21,22^. In the same surgery, microwire electrode arrays were implanted into ventral premotor cortex, i.e., part of the perilesional cortex, and a depth electrode was inserted to target subcortical motor thalamus (Fig. 1C). Post-operative CTs were fused with pre-operative CTs and MRIs to localize subcortical probe location. Location was determined to be sensorimotor thalamus for Monkey N (Fig. 1D, Fig. S2) and the internal capsule just outside sensorimotor thalamus for Monkey H (Fig. S2).

**Figure 1.**
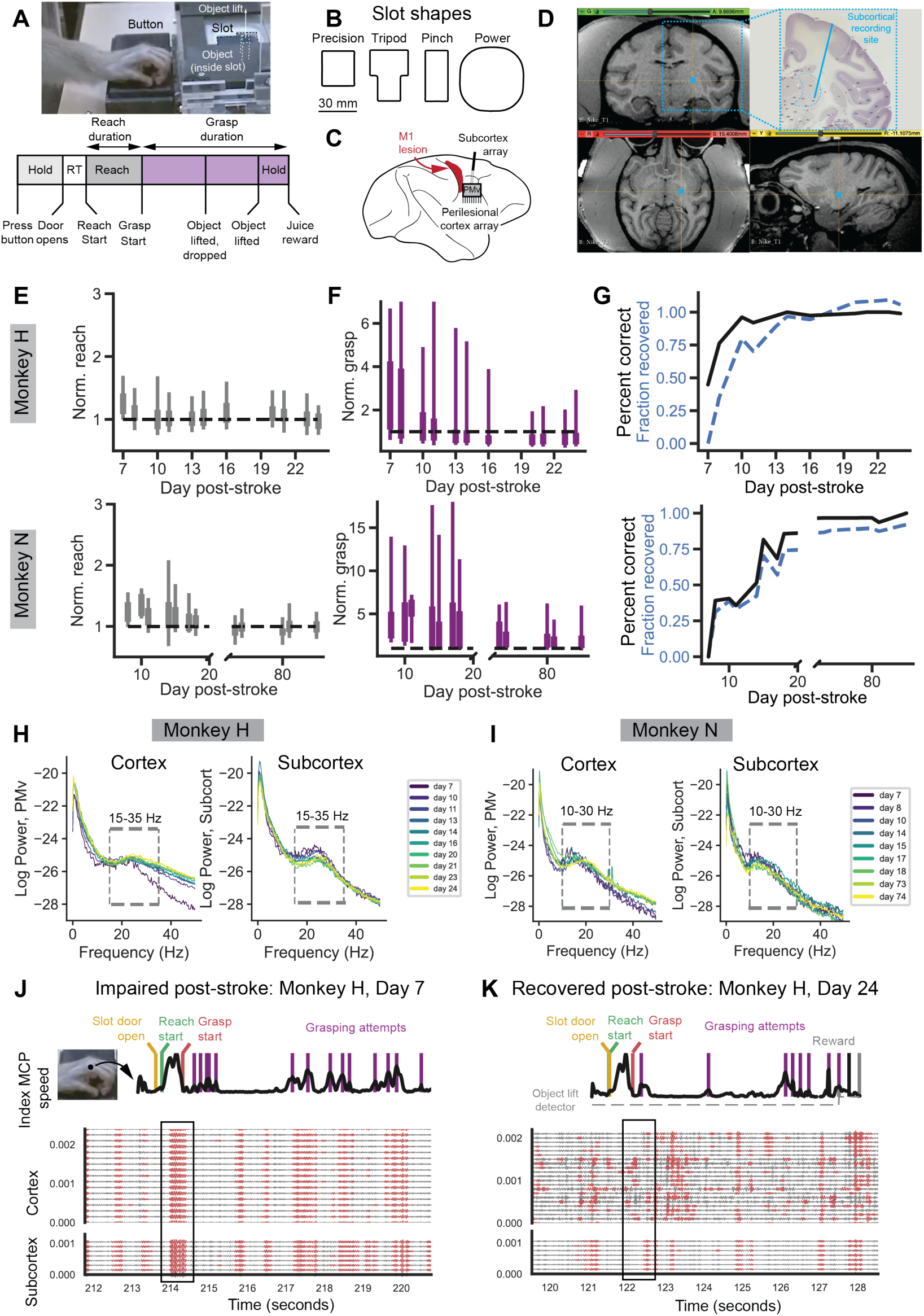
A. Reach-to-grasp task illustration and trial timeline. Animals were trained to depress a button for a specific hold time (Monkey H: 0.2 seconds, Monkey N: 1.5 seconds). Upon successfully holding, a door covering the slot and object retracted allowing access to the object. Animals reached through the slot (one of four possible slot shapes on a given trial) to grasp and lift the object. The object had to be lifted a height of approximately 1.5 cm for at least 0.267 seconds (object hold time) to trigger a juice reward. The time period from reach onset to grasp start was designated as “reach duration” and from grasp start to reward start as “grasp duration”. B. Illustration of the four different slot types that could be presented. Each was designed to elicit a unique grasp type. C. Schematic of primary motor cortex lesion and electrophysiology recordings from perilesional motor cortex and subcortex. D. T1 weighted MRI from Monkey N superimposed with subcortex probe location calculated from post- operative CT. See Supplemental Figure 1. E. Significant improvement in normalized reach duration as a function of days after stroke (linear regression, top: Monkey H, slope = -0.004, t-statistic = -2.276, N = 819, p-value = 0.023, Monkey N, slope = -0.003, t-statistics = -9.575, N = 447, p-value < 0.001) F. Significant improvement in normalized grasp duration as a function of days after stroke (linear regression, top: Monkey H, slope = -0.086, t-statistic = -7.973, N = 819, p-value < 0.001, Monkey N, slope = -0.034, t-statistics = -6.324, N = 447, p-value < 0.001) G. Significant improvement in percent correct of trials as a function of days after stroke (linear regression, top: Monkey H, slope = 0.018, t-statistic = 2.596, N = 11 days, p-value = 0.029, Monkey N, slope = 0.007, t-statistic = 4.271, N = 13 days, p-value = 0.001) H. Trial-averaged, channel-averaged power spectral densities of local field potential (LFP) signals from cortex (left) and subcortex (right) electrode arrays for Monkey H. Colors indicate different days post-stroke. I. Same H for Monkey N. J. Example trial from Monkey H, Day 7 post-stroke. Top row shows Index MCP speed (black trace) and extracted behavioral markers. Traces below show LFP channel data filtered in the beta band, with extracted beta burst epochs in red. K. Same as K, but for an example trial from Monkey H, Day 24 when the animal has recovered.

As expected, behavior improved over time as animals recovered from the stroke (Fig. 1E-G). Also as expected from prior work, there was variability in recovery timeframes despite the same lesioning approach used^21,22^. While Monkey H exhibited a typical recovery within ∼2 weeks, Monkey N took longer. He exhibited a typical, though incomplete, recovery up to day 18, after which he had an atypical worsening in behavioral performance (Fig S3). Following a period of worsening, Monkey N then exhibited a full recovery of behavior over time. In order to ensure we compared periods with an initial grasping impairment and periods with largely recovered function consistently in both animals, we chose to analyze Monkey N’s recovery up to day 18 and then compared the fully recovered days (day 73 onwards). Our choice of sessions did not significantly change the overall estimate of recovery rate in Monkey N (Fig. S3).

To study the recovery of reaching and grasping separately, trials were segmented into reaching and grasping epochs. Each trial’s reach and grasp times were normalized by the pre-stroke mean of trials to that slot such that a value of “1” indicates performance on-par with pre-stroke data. Reaching behavior, grasping behavior, and percentage of trials performed correctly significantly improved with recovery in both animals (Fig. 1E-G). To assess how neural signals change as a function of recovery, we developed a “recovery index” where a value of 0 indicated minimal recovery as recorded on the first behavioral session after the stroke and a value of 1 indicated behavior on-par with pre-stroke behavior (Fig. 1G).

### Identifying global and local beta bursts

We first assessed field potential activity from cortex and subcortex recordings (Fig. 1H,I). We consistently saw power spectrums with peaks in the beta band frequency range of ∼10-40 Hz on both the cortical and subcortical array over the course of recovery. We next studied how beta band activity was coordinated between cortex and subcortex during animal behavior. We customized beta frequency ranges for each animal based on a data-driven approach (Monkey H: 15-35 Hz, Monkey N: 10-30 Hz)^23^ and extracted beta burst events from all cortical and subcortical channels^13,16^. In general, we found that early after stroke beta burst events were strikingly coordinated both within cortex and across cortex and subcortex regions (Fig. 1J, Fig. S4, Fig. S5). Later after some recovery from stroke, we noticed that beta events were more isolated to groups of a few cortical channels and were often not as coordinated with subcortical beta bursts (Fig. 1K, Fig. S4, Fig. S5).

This observation motivated us to segment beta burst events into two different categories based on the fraction of cortical channels that were simultaneously bursting together. We observed that when pooling over behavioral sessions across recovery that there was a bimodal distribution of the fraction of cortical channels in each burst (Fig. 2A). To separate the bimodal distribution of bursts into two categories, we fit a Gaussian mixture model with two Gaussians to a session-averaged distribution. The crossing point between the two fit Gaussians was used a threshold for determining whether a particular burst event was in the category of “few isolated cortical channels bursting” or “many simultaneous cortical channels bursting”. As observed by the examples in Fig. 1JK, the fraction of burst events that were categorized as “few isolated cortical channels bursting” significantly increased with recovery from stroke (Fig. 2B). Note that our segmentation of bursts into “isolated” and “simultaneous” was based solely on cortical channel bursting, though we observed that “simultaneous” cortical bursts often coincided with subcortical bursts whereas “isolated” ones did not. We confirmed that “isolated” events were significantly less likely to co-occur with subcortical bursts compared to “simultaneous” events (Fig. 2C). For this reason, we called the “isolated” events “local beta burst events” referring to their restriction to few channels in cortex, and the “simultaneous” events as “global beta burst events” referring to their inclusion of a large fraction of cortical channels and a high likelihood of the co-occurrence of subcortical bursting.

**Figure 2.**
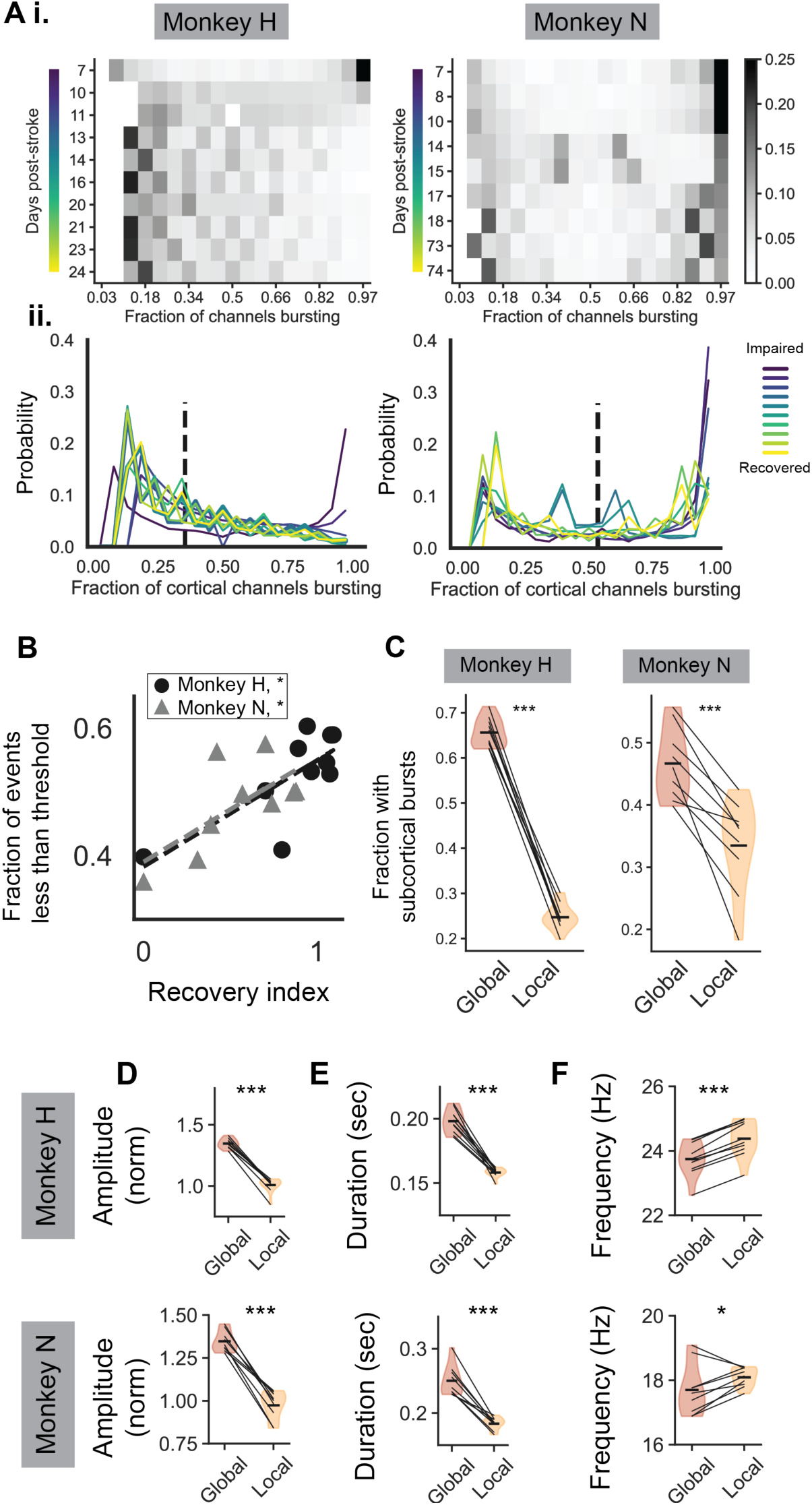
A. The distribution of the fraction of cortical channels that are bursting for each timepoint that is part of a cortical burst event. i) Each row correspond to a post-stroke day, and darker colors indicate higher occurrence. The same data is represented in (ii) as a histogram with different colors correspond to different days post stroke. The vertical dashed black line indicates the threshold calculated to separate events into local, independent burst events and global, synchronous burst events (“*local_global_threshold*”) B. The fraction of total cortical burst events that are less than the *local_global_threshold* is significantly correlated with recovery index for both animals (Monkey H: slope = 0.167, t-statistic=3.324, N = 10, p-value = 0.010, Monkey N: slope = 0.164, t-statistic = 2.393, N = 9, p-value = 0.048). C. Fraction of timepoints when cortical burst events coincide with subcortical bursting is significantly higher for global compared to local events (Monkey H: frac. timepoints during global events with subcortical bursting = 0.651, frac. timepoints during local events with subcortical bursting = 0.240, N global timepoints = 1,979,463, binomial test p-value < 0.001, Monkey N: frac. timepoints during global events with subcortical bursting = 0.459, frac. timepoints during local events with subcortical bursting = 0.361, N global timepoints = 913,342, binomial test p-value < 0.001). Error bars are s.e.m. D. Day-averaged normalized amplitudes of global bursts are significantly higher than local bursts across days (LME model with day as a random effect: Monkey H: slope = -0.169, z-statistic = - 20.296, N = 20, p < 0.001, Monkey N: slope = -0.187, z-statistic = -12.505, N = 18, p < 0.001). E. Day-averaged durations of global bursts are significantly than local bursts across days (LME with day as a random effect: Monkey H: slope = -0.020, z-statistic = -12.783, N = 20, p < 0.001, Monkey N: slope = -0.033, z-statistic = -8.590, N = 18, p < 0.001). F. Day-averaged frequency of global bursts is significantly lower than local bursts across days (LME with day as a random effect: Monkey H: slope = 0.316, z-statistic = 15.833, N = 20, p < 0.001, Monkey N: slope =0.197, z-statistic = 1.979, N = 18, p = 0.048).

Local and global beta bursts exhibited distinct burst characteristics. We found that global burst events had significantly higher amplitudes, significantly longer durations, and significantly slower frequencies than local bursts (Fig. 2D-F), suggesting that these bursts may have distinct mechanisms of generation.

One possibility we considered was that local beta bursts are just random, noisy events occurring on the cortical microelectrode array that are subthreshold global beta events. If this were the case, we would expect local bursts to involve cortical recording channels that are randomly distributed throughout the recording array (Fig. 3A, *left*). As we see in the single trial example (Fig 3B), local beta bursts tend to be more spatially clustered on the array than what would be expected by random chance (Fig. 3C) both during reaching and grasping local burst events. In addition, local bursts are significantly more spatially clustered than global beta bursts, which would not be expected if local bursts were weak, subthreshold global beta events (Fig. 3D). Overall, we find that local bursts are unlikely to be subthreshold global bursts, but are their own distinct type of beta burst that is lower amplitude, more transient, higher in frequency, and more spatially clustered than global beta bursts (Fig. 2DEF, Fig 3CD). Below we further explore how local and global bursts differ in their task occurrence and in their relationship to spiking neural dynamics.

**Figure 3.**
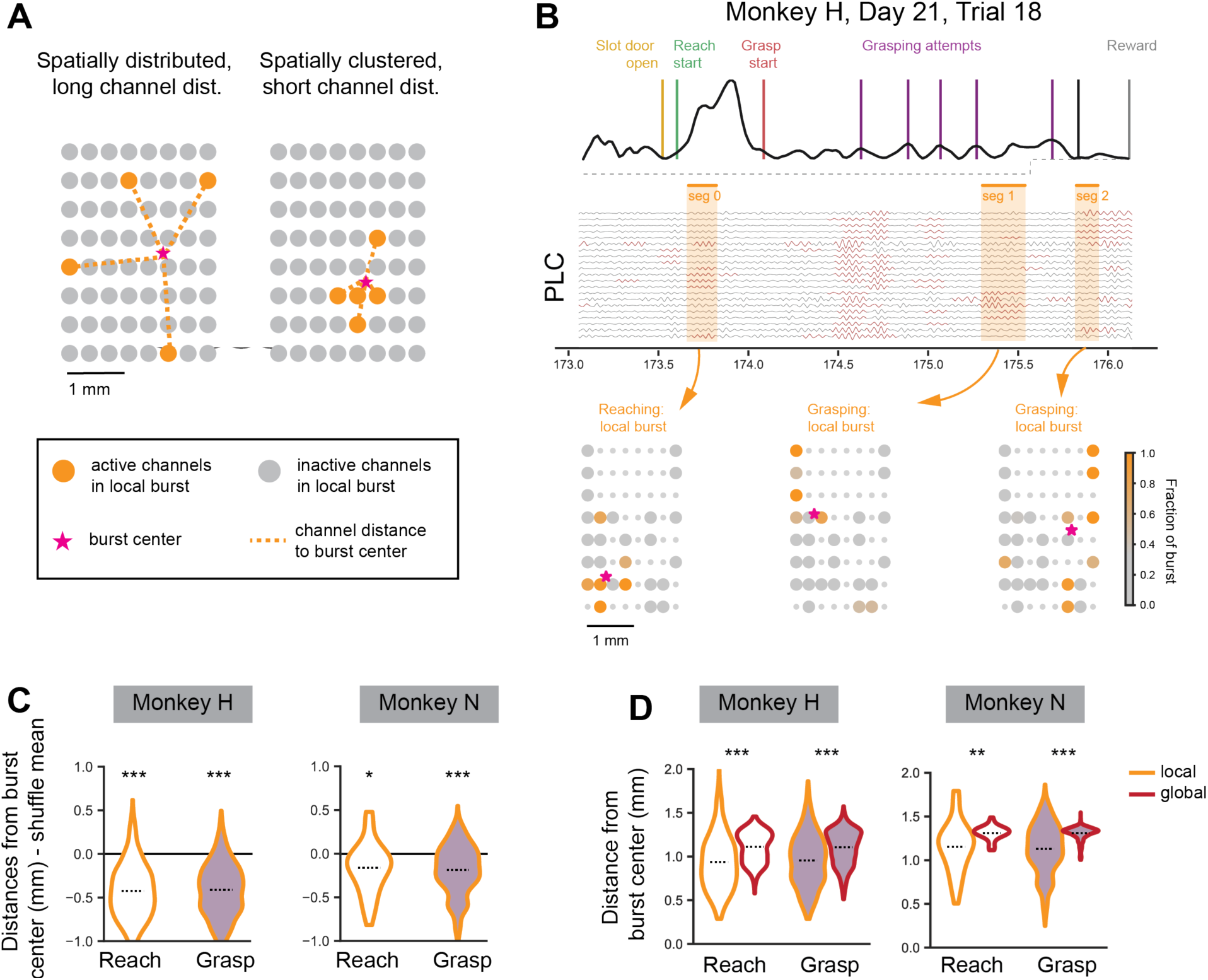
A. Example of possible spatial configurations of local beta bursts. *Left* illustrates spatially distributed burst, where the distances between channels active in the burst and spatial burst center are large. *Right* illustrates spatially clustered burst where the distances between channels active in the burst and spatial burst center are small. B. Example of the spatial distribution of active channels from local bursts on a single trial (Monkey H, day 21, trial 18). Color of each channel indicates for what fraction of the burst channels are active. Channels with small markers have been excluded due to noise. C. Individual local burst events have channel distances that are more spatially clustered than expected just based on the number and location of channels that are generally activate across all reach and grasp local burst events (LME model with day as a random effect, testing for mean of distribution being different than zero: *left*: Monkey H, *reach*: intercept = -0.426, z-statistic = -15.590, N = 421, pv < 0.001, *grasp:* intercept = -0.407, z-statistic = -14.200, N = 1745, pv < 0.001, *right:* Monkey N: *reach:* intercept = -0.135, z-statistic = -2.116, pv = 0.034, *grasp:* intercept = -0.185, z-statistic = - 5.335, N = 1179, pv < 0.001). D. Local burst events have channel distances that are more spatially clustered than global burst events (LME model with day as a random effect: *left:* Monkey H, *reach:* slope = 0.222, z-statistic = 11.787, N = 750, pv < 0.001, *grasp*: slope = 0.178, z-statistic = 22.929, N = 3452, pv < 0.001, *right:* Monkey N, *reach:* slope = 0.0978, z-statistic = 2.587, N = 115, pv = 0.010, *grasp:* slope = 0.162, z- statistic = 21.266, N = 2566, pv < 0.001).

### Global but not local beta bursts are associated with slow movement

Global and local burst events occurred during different phases of the task and showed different dynamics with recovery from stroke. Early in stroke when animals were the most impaired, global beta bursts occurred frequently around grasp start (red box Fig. 4A, Fig. S6A). Grasp start is a behavioral marker that indicates the time point at which the animals have completed transporting their hand towards the slot and the grasping phase of the task is beginning. This occurrence of global beta bursts around grasp start significantly reduced with recovery (Fig. 4C). Global beta burst events did not exhibit consistent significant changes when aligned to other task behavioral events. In contrast, local beta events did not show significant changes aligned to particular behavioral events, but did show significant increases throughout the trial period spanning from reach start to juice reward (red box Fig. 4B, Fig. S6B, Fig. 4D).

**Figure 4.**
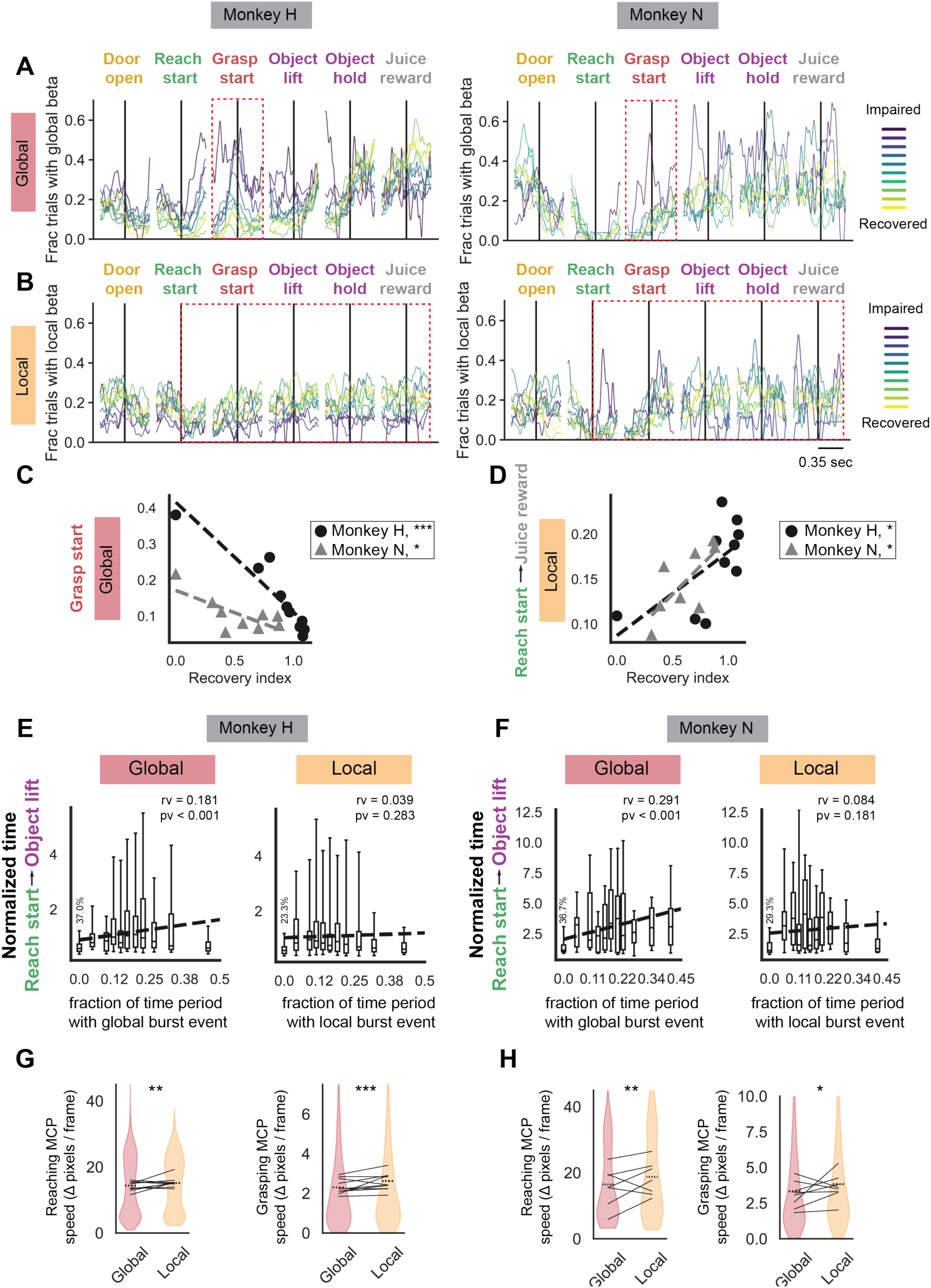
A. The fraction of trials with global beta burst events for each day post-stroke (colored trace) aligned to different behavioral events (x-axis, window of [-0.35, 0.35] seconds around each event). Red box indicates area of notable change in rate of burst events with recovery. B. Same as (A) but for local beta bursts events. C. Recovery index is significantly negatively correlated with the mean fraction of trials with global beta events in a [-0.35, 0.35] second window centered at grasp start (Monkey H: slope = -0.305, t-statistic = -7.364, N = 10, p-value < 0.001, Monkey N: slope = -0.123, t-statistic = -2.798, N = 9, p-value = 0.027). Days with LFP and behavior are included. D. Recovery index is significantly positively correlated with the mean fraction of trials with local beta events in the reach start to juice reward (Monkey H: slope = 0.093, t-statistic = 2.340, N = 10, p-value = 0.047, Monkey N: slope = 0.125, t-statistic = 2.546, N = 8, p-value = 0.044). Days with LFP and rewarded trials are included. E. There is a significant positive correlation between time between time spent in a global beta burst event and trial length (*left:* Monkey H: slope = 1.47, t-statistic = 5.096, N = 765 trials, p-value < 0.001), but no significant correlation between time spent in a local beta burst event and trial length (*right:* Monkey H: slope = 0.34, t-statistic = 1.075, N = 765 trials, p-value = 0.283). Regression is performed on all trials pooled. Data is visualized in percentiles: Trials with no beta burst events are all in first boxplot aligned with “0.0” on the x-axis and percentage of trials is indicated in text above the box plot. Remaining trials are grouped into 10 percentiles based on fraction of time spent in burst event that correspond to the 10 boxplots to the right of the first boxplot. Percentiles are for visualization only, statistics are performed on individual trials. F. Same as (E) for Monkey N (*left:* slope = 5.50, t-statistic = 4.839, N = 256 trials, p-value < 0.001, *right*: slope = 1.69, t-statistic = 1.343, N = 256 trials, p-value 0.181) G. Hand speed (measure with index finger MCP speed) is significantly faster during local beta burst events than during global beta burst events for both reaching (reach start to grasp start) and grasping epochs (grasp start to last object lift) (*left*: reaching, LME with day as random effect: slope = 0.934, z-statistic = 3.167, N = 3828 camera frames, p-value = 0.002, *right:* grasping, LME with day as random effect: slope = 0.259, z-statistic = 3.697, N = 6144 camera frames, p-value < 0.001). Error bars are s.e.m. H. Same as (G) for Monkey N (*left*: reaching, LME with day as random effect: slope = 3.379, z-statistic = 2.781, N = 408 camera frames, p-value = 0.005, *right:* grasping, LME with day as random effect: slope = 0.276, z-statistic = 2.023, N = 2365 camera frames, p-value = 0.043) Error bars are s.e.m.

We can gain more insight into the relationship between beta burst events and behavior at the single-trial level. In single trials, we found that the fraction of the time between reach start and last object lift that is part of a global burst event was significantly correlated with the amount of time that a trial takes (Fig. 4E-F, *left*). In other words, when trials were dominated by global burst events, they took significantly longer to complete. This relationship did not hold up for local beta burst events (Fig. 4E-F, *right*). This single trial analysis implies that global beta burst events reflect slowed movement periods. We confirmed that hand speed was significantly slower during global bursts events than local burst events for both the reaching epoch (Fig. 4G-H, *left*) and grasping epoch (Fig. 4G-H, *right*). Overall, we found that global beta events occurred early after stroke during grasping periods and were associated with slower trial times and slower reach and grasp movements. On the other hand, local beta events emerged during movement as animals recovered from stroke but were not significantly linked to trial duration.

### Recovery is associated with increases in grasp-epoch firing rate and firing variation

We next sought to assess how local and global beta bursts interacted with single unit and population level firing properties over the course of recovery. Single units were detected on individual behavioral sessions and were analyzed as a group for changes in reaching and grasping epoch firing properties over the course of recovery. Due to poor unit yield in sessions early after stroke in one animal (monkey H) and diminishing unit yield in sessions with recovery in the second animal (monkey N), we divided sessions into “early” and “late” categories to pool units across sessions for each animal for a more balanced comparison (see Table S1, Fig. S7). Representative trial-averaged z-scored single unit patterns aligned to reach start (Fig. 5A) and grasp start (Fig. 5B) show the general trend that reach-related single units are relatively equally modulated during early and late sessions whereas grasp-related single units show increases in modulation with recovery (Fig. 5AB, *bottom*). This was not surprising given that animals exhibited mild reaching deficits that recovered relatively quickly in comparison to animals’ more severe grasping deficits (Fig. 1E-F). To quantify this observation, we measured each recorded units’ firing rate and firing rate variation during reaching epochs and grasping epochs. We found that there were no significant differences in mean firing rate and firing rate variation in units during reaching epochs from early to late sessions (Fig. 5C-D, *gray*), but there were significant increases in mean firing rate and firing rate variation in units during grasping epochs from early to late sessions (Fig. 5C-D, *purple*). Overall, units significantly increased their grasp-epoch firing rates and firing rate variation from early to late recovery sessions.

**Figure 5.**
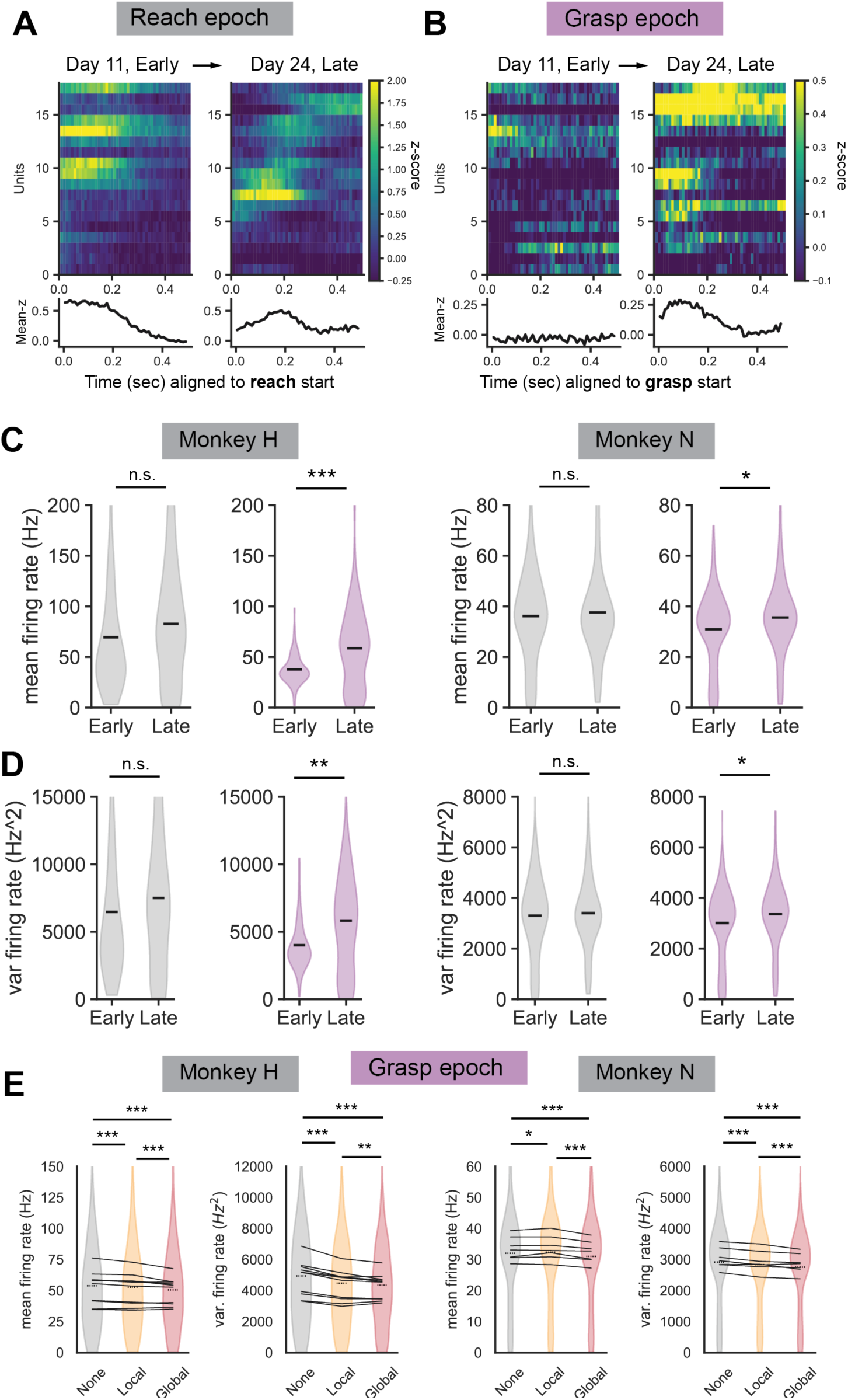
A. Example of trial-averaged z-scored spiking activity spiking units from day 11 and day 24 aligned to reach start (Monkey H). Bottom shows average z-scored activity across all displayed units. B. Same data as (A) but aligned to grasp start. C. There is no significant change in trial-averaged firing rates of units during reaching from early sessions versus late sessions (independent sample t-test*, left gray:* Monkey H: t-statistic: 1.633, N = 279, p-value = 0.104, *right gray:* Monkey N: t-statistic = 0.595, N = 339 units, p-value = 0.552), but there is a significant increase in trial-averaged firing rates during grasping from early sessions versus late sessions (independent sample t-test, *left purple:* Monkey H: t-statistic: 4.067, N = 279, p-value < 0.001, *right purple:* Monkey N: t-statistic: 2.466, N = 339, p-value = 0.014). Bars are means and s.e.m. D. There is no significant change in trial-averaged firing variability of units during reaching from early sessions versus late sessions (independent sample t-test*, left gray:* Monkey H: t-statistic: 1.381, N = 279, p-value = 0.168, *right gray:* Monkey N: t-statistic = 0.503, N = 339 units, p-value = 0.615), but there is a significant increase in trial-averaged firing variability during grasping from early sessions versus late sessions (independent sample t-test, *left purple:* Monkey H: t-statistic: 3.412, N = 279, p-value = 0.001, *right purple:* Monkey N: t-statistic: 2.044, N = 339, p-value = 0.042). Bars are means and s.e.m. E. There are significant differences between the trial-averaged mean firing rate and variability of firing rate of units during grasping when there are no beta events (gray), local beta events (yellow), and global beta events (red). (paired t-test, *left:* Monkey H, all N = 279 units, *mean firing rate:* None vs. local: t-statistic = 6.333, p-value < 0.001, None vs. global: t-statistic = 6.969, p-value < 0.001, Local vs. global: t-statistic = 4.535, p-value < 0.001, *firing rate variability:* None vs. local: t-statistic = 13.141, p-value < 0.001, None vs. global: t-statistic = 10.329, p-value < 0.001, Local vs. global: t-statistic = 3.464, p-value = 0.002, *right:* Monkey N, all N = 339 units, *mean firing rate:* None vs. local: t-statistic = -2.547, p-value = 0.034, None vs. global: t-statistic = 7.081, p-value < 0.001, Local vs. global: t- statistic = 7.812, p-value < 0.001, *firing rate variability:* None vs. local: t-statistic = 5.449, p-value < 0.001, None vs. global: t-statistic = 11.354, p-value < 0.001, Local vs. global: t-statistic = 4.274, p-value < 0.001). All p-values are Bonferroni corrected for multiple comparisons.

### Entrainment of single units to global bursts slow firing rate and firing variation

Given the changes in grasp-epoch firing properties of units, we next focused on how the presence of global and local beta burst events influenced grasp-epoch firing properties of units. We found that the grasp-epoch mean firing rate and firing rate variability were significantly lower during global burst events compared to no beta bursting events and compared to local burst events (Fig. 5E). Firing variability was significantly lower during local burst events compared to no burst events, while firing rate was lower in one animal (Monkey H) and higher in the second animal (Monkey N) during local burst events compared to no burst events.

To link beta bursts more tightly to reductions in firing rate and firing variation, we sought to quantify the influence of beta bursts on individual units through measurements of phase entrainment. Example single unit waveforms and entrainment to local and global beta burst events are illustrated in Fig. 6A. Entrainment is quantified by calculating each units’ phase histogram resultant vector length (*r*), and normalizing this by the mean and standard deviation of a shuffle distribution of *r* for that unit yielding a “modulation z-score”. In the representative units, the top unit shows stronger entrainment to global beta events whereas the bottom unit shows stronger entrainment to local beta events. Although individual units occasionally exhibit stronger entrainment to local beta events than to global beta events, the overall modulation z-scores of significantly entrained units (z-score > 95th percentile of the shuffle distribution) are significantly higher for global beta bursts compared to local beta bursts (Fig. 6B).

**Figure 6.**
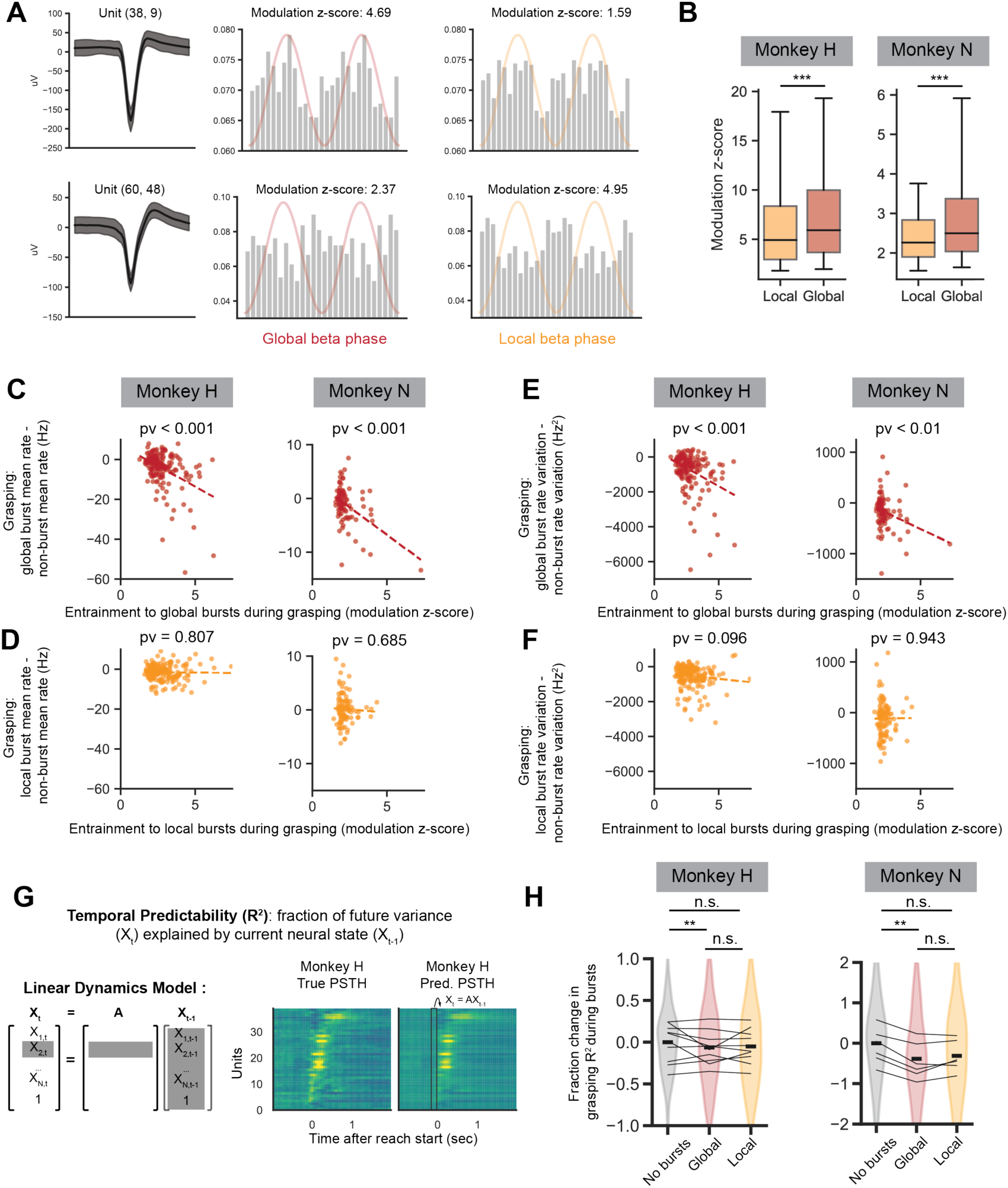
A. Examples of isolated single unit waveform mean +/- 1 standard deviation (left), unit entrainment to global beta events (middle), and local beta events (right). Probability of unit firing is plotted against global and local beta phases. Example units are from Monkey H, day 24. B. Distribution of modulation z-score values for significantly entrained unit-channel pairs. Units are more strongly entrained to global beta bursts than local beta bursts (independent sample t-test: *left*: Monkey H: t-statistic: 8.096, N = 9872 significant unit-channels, p-value < 0.001, *right:* Monkey N: t-statistic: 12.879, N = 5779 unit-channels, p-value < 0.001). Bars span 5^th^ to 95^th^ percentile of data. C. There is a significant relationship between entrainment to global beta bursts during grasping (modulation z-score) and change in trial-averaged mean firing rate during grasping between global burst times and non-burst times (linear regression, *left:* Monkey H: t-statistic = -5.960, N = 174, p- value < 0.001, *right*: Monkey N: t-statistic = -4.856, N = 90, p-value < 0.001). D. There is no significant relationship between entrainment to local beta bursts during grasping (modulation z-score) and change in trial-averaged mean firing rate during grasping between local burst times and non-burst times (linear regression, *left:* Monkey H: t-statistic = -0.245, N = 200, p- value = 0.807, *right*: Monkey N: t-statistic = -0.407, N = 104, p-value = 0.685). E. There is a significant relationship between entrainment to global beta bursts during grasping (modulation z-score) and change in trial-averaged firing rate variation during grasping between global burst times and non-burst times (linear regression, *left:* Monkey H: t-statistic = -4.837, N = 174, p-value < 0.001, *right*: Monkey N: t-statistic = -2.878, N = 90, p-value = 0.005). F. There is no significant relationship between entrainment to local beta bursts during grasping (modulation z-score) and change in trial-averaged firing rate variation during grasping between local burst times and non-burst times (linear regression, *left:* Monkey H: t-statistic = -1.671, N = 200, p-value = 0.096, *right*: Monkey N: t-statistic = 0.071, N = 104, p-value = 0.943). G. Schematic of neural population dynamics model used to assess temporal predictability. H. There is a significant fraction change in grasping temporal predictability (population dynamics model R^2^) during global (red) but not local (yellow) beta bursts (LME with day as random effect: *left*: Monkey H, *none vs. global*: slope = -0.018, z-statistic = -2.825, N = 1323 valid trials, p-value = 0.0142, *none vs. local:* slope = -0.004, z-statistic = -2.015, N = 1396 valid trials, p-value = 0.132, *global vs. local*: slope = 0.003, z-statistic = 0.692, N = 1151 valid trials, p-value = 1.0, *right:* Monkey N, *none vs. global:* slope = -0.027, z-statistic = -3.145, N=363, p-value = 0.005, *none vs. local*: slope = -0.007, z-statistic = -2.187, N = 371, p-value = 0.0863, *local vs. global:* slope = 0.003, z- statistic = 0.605, N = 322, p-value = 1.0). All p-values are Bonferroni corrected for multiple comparisons. Bars are mean +/- s.e.m.

Finally, we link each units’ entrainment to global and local burst events during grasp epochs to their change in firing rate and firing rate variation during grasp epochs. We find that stronger entrainment to global beta events predicts greater reductions in firing rate (Fig. 6C) and firing variability (Fig. 6E), however, that entrainment to local beta events do not show these trends (Fig. 6E, F).

Altogether, both global and local beta bursts entrain units during the grasp epoch. Entrainment to global bursts but not local bursts is significantly correlated with reductions in firing rate and firing variability that might impede the increases in grasp-epoch firing rate and variability needed for grasp execution. Entrainment to local bursts does not significantly predict changes in firing rate or variability, suggesting that local bursts may reflect a fundamentally different neural computation than global bursts.

### Interruption of population neural dynamics during global but not local beta bursts

In our previous work, we established that spatiotemporal patterns of neural populations become more temporally predictable as animals recover from stroke^21^. Specifically, when fitting a linear model that predicts the future neural population activity pattern from the current pattern (Fig. 6G), the predictability of the model has been shown to increase as animals recover, supporting that population neural patterns become more stereotyped with recovery. Here, we ask how the occurrence of local and global beta events influence the predictability of neural patterns. True and model-predicted population spiking patterns during grasping periods were collated during non-bursting bins, global bursting bins, and local bursting bins. The variance explained by the model-predicted bins was calculated for all three groups (Fig. 6H). Variance explained by model-predicted bins occurring during global bursts was significantly lower than variance explained by non-bursting bins. Variance explained by local bins was not significantly different from non-bursting bins or global bursting bins. Overall, this supports that predictable spatiotemporal patterns of spiking that have been shown to become more predictable with recovery are interrupted during global but not local bursts.

### Local and global beta events are separable in neurotypical primates

While our previous analysis focused on primates recovering from motor cortex stroke, we also find separable local and global beta events in intact animals. Two separate animals were implanted with microelectrode arrays targeting primary motor cortex (M1) and trained to execute a sequential target-reaching task (Fig. 7A). Notably, this task requires rapid planning of future movements, even when a current movement is still being finished. This design is more complex than prior studies investigating beta dynamics in simple movements, where preparation and execution are typically separated in time.

**Figure 7.**
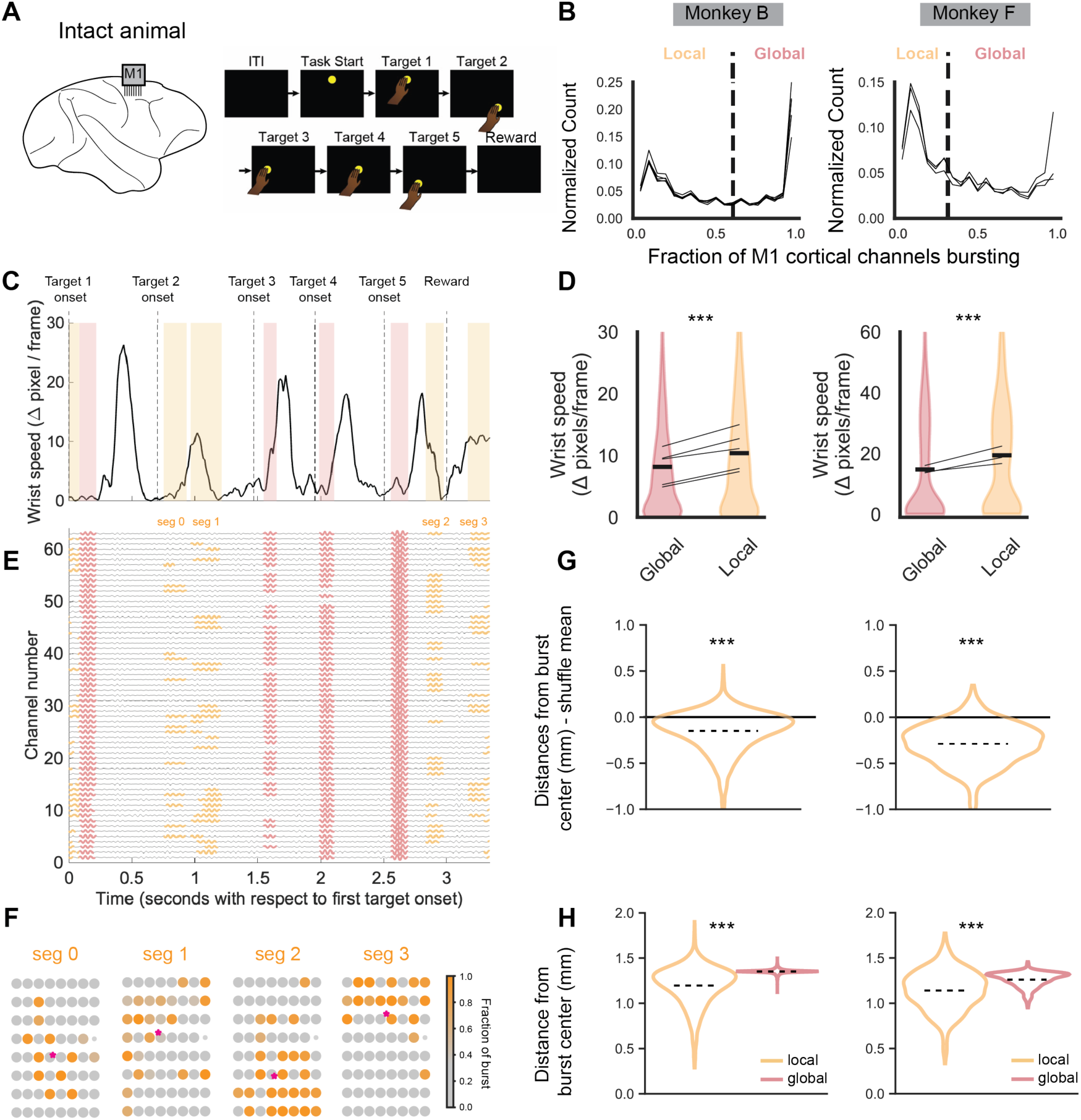
A) Neural recordings were obtained from primary motor cortex in animals trained to perform a sequential target-capture task. B) Distribution of fraction of channels bursting simultaneously during beta bursts. Different lines indicate different behavioral days for each animal. Thresholds used to separate local and global beta events are indicated with horizontal dashed lines. C) Single trial wrist speed with global and local events highlighted using red and yellow boxes respectively (trial: Monkey B, session 0, trial 55). D) Wrist speed is significantly lower during global versus local beta events (LME with day as random effect: *left* Monkey B: slope = 2.63, t-statistic = 11.40, N = 7136, p < 0.001, *right* Monkey F: slope = 4.61, t-statistic = 5.46, N = 1790, p < 0.001). Bars are mean +/- s.e.m. E) Channel level activity filtered in the beta band during single trial shown in (C). Global and local burst events are shown in red and yellow respectively. F) Example of the spatial distribution of active channels from local bursts on a single trial shown in (C, E). G) Individual local burst events have channel distances that are more spatially clustered than expected just based on the number and location of channels that are generally activate across all reach local burst events (LME model with day as a random effect, testing for mean of distribution being different than zero: *left*: Monkey B, intercept = -0.148, z-statistic = -10.32, N = 3557, pv < 0.001, *right* Monkey F, intercept = -0.304, z-statistic = -10.917, N = 867, pv < 0.001). H) Local burst events have channel distances that are more spatially clustered than global burst events (LME model with day as a random effect: *left:* Monkey B, slope = 0.158, z-statistic = 43.53, N = 7173, pv < 0.001, Monkey F, slope = 0.122 z-statistic = 15.83, N = 1824, pv < 0.001)

When assessing the fraction of cortical channels simultaneously bursting, there was a clear bimodal distribution in both animals (Fig. 7B). We used the same Gaussian mixture model approach to identify a threshold between local and global events. We found that global beta events tended to occur during non-movement periods between reaches whereas local beta events could occur during movements if they were low speed (Fig. 7C). As in the stroke recovery data, hand speeds were significantly lower during global beta bursts compared to local beta bursts (Fig. 7D). We also confirmed in the healthy animals that local beta events were significantly more spatially clustered than expected given random chance (Fig. 7F-G) and more spatially clustered than global beta bursts (Fig. 7H). Overall, we find that behavioral and spatial clustering characteristics of local and global bursts identified in stroke animals also hold up in motor cortical regions of neurotypical animals.

## Discussion

Overall, in stroke and neurotypical non-human primates performing complex behaviors, we find that beta burst events are dissociable into local events that are limited to few spatially clustered channels in cortex and global events that involve many simultaneously bursting cortical channels that tend to co-occur with subcortical beta bursts. Global bursts occur more frequently in movement-impaired animals, occur mid-movement, and are associated with slow movement speeds. On the other hand, local bursts occur more frequently in recovered animals throughout behavior. Fig. 8 summarizes these findings.

**Figure 8.**
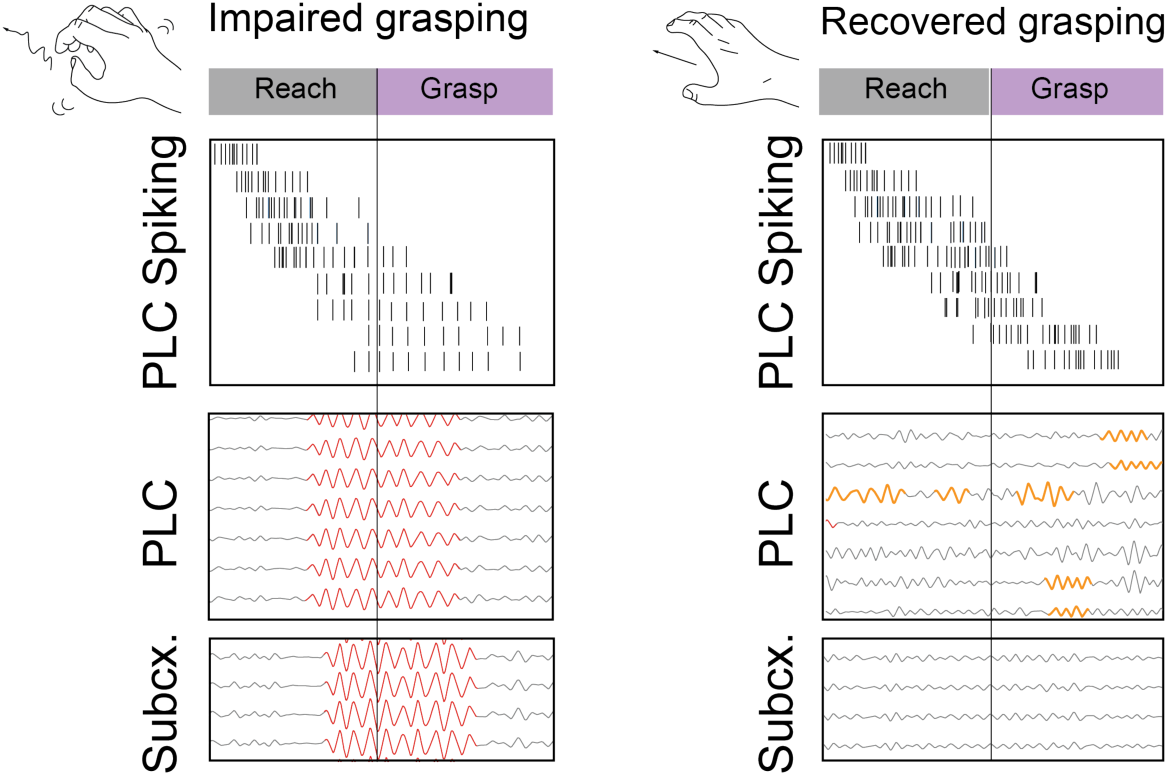
Summary of main findings. In impaired animals (*left)*, global bursts (red) that are synchronous across perilesional cortex (PLC) and subcortex (Subcx) occur mid-movement and are associated with slower movement speeds, longer trial times, entrainment of single unit activity, a reduction in single unit firing rate and firing variance, and interrupted spiking population dynamics. In recovered animals (*right*), global bursts during movement are uncommon but local bursts (orange) occur during movement and are associated with the recovered state, typical single unit firing rate and firing variance, and typical spiking population dynamics.

### Dichotomy of global versus local beta bursts

The identification of separate global and local beta bursts required a focus on beta burst events^13,16^ instead of continuous, channel-averaged beta power. If we had only analyzed channel-averaged beta power, we would have missed the local beta bursts due to their lower amplitude and fewer number of involved channels. This finding also required an electrophysiology preparation with a cortical multi-electrode array and a subcortical recording probe. Multiple contacts in a single cortical area were required for detecting synchronous beta bursts, and the subcortical probe was required to associate the synchronous bursts with subcortical bursts. Previous studies in humans undergoing deep brain stimulation surgery or with a chronically implanted DBS device plus cortical ECoG strip have such a preparation^24^, but the density of recording sites in a single cortical area is limited. In non-human primate movement control research, there are few labs that simultaneously record from cortical and subcortical sites. A further impediment to the observation of global versus local beta bursts may be certain referencing schemes used to pre-process the LFP signal -- bipolar signal acquisition, or common average referencing on a cortical array would subtract away substantial segments of the global beta oscillations.

While the specific dichotomy of global and local beta bursts has not been previously reported, other studies point towards the findings of synchronous and isolated beta bursts in motor cortex. A recent study from Denker et. al.^25^ focuses on beta phase patterns across an intracortical array during movement preparation. While the focus of their study is on their findings of traveling waves in planar, circular, radial, and rotating configurations, they also find synchronous and random phase patterns. Since their study does not use a beta amplitude threshold to determine which channels are engaged in bursting, it is not directly comparable to the classification scheme in our study. However, based on when the different spatiotemporal patterns tend to occur during a reach-to-grasp behavior^25^, it seems likely that the synchronous waves correspond to global beta events as they tend to both occur prior to and following movement. The random waves likely correspond to local beta events as they both tend to occur throughout behavior, especially during grasping movements. Note here that “random” refers to random beta phase patterns, not random channels.

### Global bursts and slowed movement

Here, the occurrence of global beta oscillations corresponded to periods of slowed movement in both stroke-impaired and intact animals. Given that global beta oscillations tend to co-occur with subcortical beta oscillations, this observation is consistent with the Parkinson’s disease literature that finds increased subcortical beta in PD movement-impaired patients compared to non-PD patients^26,27^. We also find that global beta oscillations entrain cortical neurons, consistent with findings in PD patients that the amplitude of high gamma signals, a possible proxy for multi-unit spiking activity, in sensorimotor cortical ECoG phase-lock to beta much more than in non-PD patients ^26,28–30^. Here we find that the degree of entrainment to global beta predicts how strongly neural activity firing rates and firing rate variability will decrease during global beta events. Since recovery from stroke is associated with increased firing rate modulation around grasp periods (Fig 5CD), this evidence supports the possibility that over-entrainment of cortical neurons to global beta impairs the neural modulation needed for effectively generating signals for control of movement. This phenomenon is consistent with theories of how exaggerated subcortical beta in PD disrupts cortical neural patterns needed to generate movement^28^. Lastly, we find that global beta bursts that seem to impair movement are longer duration than local beta bursts (Fig 2E), consistent with the finding in PD that movement impairment is associated with longer duration beta bursts^31^. More recent work has designed adaptive deep brain stimulation in PD patients to shorten beta burst duration and has demonstrated clinical improvement greater than conventional deep brain stimulation^32^. This is suggestive of an intervention post-stroke that specifically targets long-duration global beta bursts.

### Mechanisms resulting in excessive global beta after stroke

Overall, we find high global beta early in stroke when animals exhibit movement impairment. In PD, a lack of dopamine in the striatum is thought to compromise its ability to attenuate beta oscillations, resulting in the excessive subcortical beta observed in PD^28,33^. However, in stroke, the origin of the excessive global beta is less clear. One candidate mechanism is the known increase in GABA following an infarct^34^. Increases in GABAA-currents have been demonstrated to increase the power of beta oscillations and slightly reduce their peak frequency^18^. In our study, we do observe an increase in the peak frequency of beta with recovery (Fig.1 HI), possibly mirroring the reduction in excess GABA. Another possible mechanism is related to the commonly observed post-stroke increase in resting state low frequency oscillations in cortex and subcortex^35–37^. Low frequency oscillations are thought to emerge due to the disconnection of cortex from subcortical nodes, a phenomenon that drives subcortex into an autonomous oscillatory mode^36,38^. A recent neural network model of cortical beta oscillations demonstrates that external inputs into cortex that have a high autocorrelation (i.e. are low frequency) tends to drive longer duration beta bursts^39^ (Fig. 2, Supp. 3, Panel G, P). Thus, it is possible that the exacerbated post-stroke non-task related low frequency oscillations may contribute to driving intermittent global beta bursts.

Another possibility emerges from the view that beta oscillations in cortex, whether local or global, reflect an output from the cortico-basal ganglia-thalamic loop that helps stabilize cortical patterning. In the neurologically-intact brain, coordinated cortical co-firing patterns spatiotemporally modulate the putamen, a key basal ganglia structure involved in motor control^40^. Coordinated cortical-putamen patterns emerge with skill learning^41^, and are filtered through the rest of basal ganglia and thalamus, and conveyed back to cortex^42^. Beta oscillations output from the basal ganglia have been hypothesized to contribute to the inhibition of unselected actions or effectors^43^, or the stabilization of motor plans to prevent interference from competing actions or other salient cues^44^. For precise reach-to-grasp behaviors, transient, spatially specific local beta bursts may then reflect a finely tuned circuit that enables accurate grasping in response to sensory feedback while stabilizing the proximal arm.

After an M1 stroke, the loss of cortical-striatal coordination may lead to the breakdown of the network’s ability to output fine-tuned, localized beta bursting and result in the emergence of more global, less specific beta bursts. Recovery of skilled prehension in rodents is associated with the restoration of coordinated activity between the perilesional cortex and striatum^45^. Perhaps consistent with this model, we find that increased grasp-related firing and neural variability in ventral premotor cortex is associated recovery of prehension control and more localized beta bursts.

### Global versus local beta bursts during grasping

Why global beta oscillations tended to occur at grasp start could be due to a few different reasons. First, spiking modulation around grasp start was very weak early after stroke (Fig. 5B) This observation is corroborated by other studies^21,46,47^. It has been hypothesized that strong beta signals tend to occur during periods of weak spike co-firing^46^. Thus, the unusually weak neural co-firing at grasp start could allow global beta signals to intrude. A second possibility is that, after stroke, movements have been observed to be more fragmented and less smoothly executed^48–50^. As the reach-to-grasp task consists of a reaching movement to transport the hand, followed by a grasp, it is possible that the movement is being segmented into two separate “reach” and “grasp” movements. The global beta observed at grasp start may then just be the classic post movement beta resynchronization following the reaching component. With recovery then, the two movements may have been fused back into a single reach-to-grasp movement.

In contrast to global beta, a spatially clustered, lower amplitude, higher frequency, local-to-cortex beta was characterized. Local beta increased in prevalence with recovery from stroke (Fig. 4D), and tended to occur during grasping phases of the reach-to-grasp movement (Fig. 4B). This finding contrasts with the general notion that beta bursts are not present during movement. However, there are exceptions to this rule: beta bursting is high in cortex during object grasps when animals must output non-zero muscular force^15,51,52^, as well as during tactile exploration^53^. This reach-to-grasp task was indeed more of a tactile exploration task since animals had to reach into a slot to retrieve an object that they might not always be able to view. Multiple attempts to grasp the object were often needed. Previous work has demonstrated that neurons in the grasping network, involving parietal cortex, premotor cortex, and primary motor cortex have been shown to communicate at beta frequencies during reach-to-grasp tasks^54^. Somatosensory and motor cortices have also been shown to communicate at beta frequencies during object manipulation tasks^52^. Within premotor cortex, previous work has distinguished between low frequency beta oscillations that are anti-correlated with movement such that more low frequency beta delays movement onset, and higher frequency beta oscillations that occur prior to fast movement onset times^55^. As our local beta bursts are higher frequency than the global beta bursts (Fig. 2F), it is possible they may reflect this pro-kinetic signal.

We hypothesize that since the grasping behavior in this study consisted of multiple grasping segments such as object contact, object pinch, re-attempted object pinches, object lift, and object hold, local beta oscillations could occur intermittently during the pinch and hold phases of this task. In addition, it is possible that beta occurring in between grasping movement phases potentially reflects future grasp plans^56^. Finally, the spatial focality of the local beta bursts suggests that these bursts may encode information in a somatotopically relevant manner. Indeed, it has been shown that focal stimulation in somatosensory cortex will drive a somatotopically corresponding site in motor cortex^57,58^. Thus, particular somatosensory events experienced by specific fingers such as object contact, object slip, or object holds may drive perilesional motor cortex beta oscillations at sites that are somatotopically matched. Further adding to this notion is evidence that somatosensory stimulation can drive beta desynchronizations and subsequent synchronizations, and that the strength of these synchronizations increase with recovery from stroke^59,60^. In summary, local beta may reflect grasp planning inter-movement epochs, object pinch and/or hold epochs, or responses to salient somatosensory feedback.

### Biomarker of impairment

Previous studies have developed biomarkers of movement impairment based on beta desynchronization and resynchronization dynamics during simple movements or beta levels at rest. For example, a lack of full cortical beta desynchronization characterizes impaired movement following stroke^11^. Excessive subcortical beta power^61^, excessive coupling between high frequency signals and beta signals in cortex^28^, and excessive cortico-cortical coherence^62^ predict an impaired state in PD. Overall, our findings that excessive global beta during movement predict slower movement speeds and greater movement impairment are consistent with trends observed in previous stroke studies and Parkinson’s disease literature.

What is particularly novel in this study is that we find *increased* prevalence of local beta events during movement as animals recovered from stroke and showed behavioral improvement. As described above, we hypothesize these increases could reflect planning, object holding/pinching, or beta rebounds following the communication of salient somatosensory feedback. It is possible then, that detection of local beta during grasping may serve as an additional biomarker of sensorimotor function. While global beta may reflect a brain state that impedes movement execution, perhaps due to excess synchrony, in the cortico-basal ganglia-thalamic loop, local beta may reflect cortico-cortical communication necessary for execution of complex movements that require planning and corrections based on feedback. As suggested previously^59,60^, in order for motor area beta oscillations to be successfully driven by other cortical areas, cortical network connectivity may need to recover. The increased prevalence of local beta events following recovery of complex movement could then reflect progress of this cortico-cortical reconnection.

Overall, we were able to discover distinct categories of global and local beta burst dynamics during complex behavior in movement-impaired and neurologically intact non-human primates. Global bursts are largely consistent with previous reports of beta as an anti-movement signal, while local bursts are a novel finding that may reflect cortico-cortical communication needed for complex movement. Future work will determine the utility of separately tracking global and local signals in movement disorders.

## Acknowledgements

We thank M.J. Lemoy and K. Stotts for surgical assistance. This research was funded by the NINDS of the NIH through postdoctoral career transition award 1K99NS124748 to P.K. Research reported in this publication was supported by the NINDS of the NIH (R01NS117406, R01HD111562 to K.G.). The content is solely the responsibility of the authors and does not necessarily represent the official views of the NIH.

## Author contributions

**PK**: Conceptualization, Methodology, Software, Formal analysis, Investigation, Data Curation, Writing - Original Draft, Writing - Review & Editing, Visualization, Funding acquisition, **BF**: Formal analysis, Visualization, **HC**: Investigation, Data Curation, **SG**: Investigation, Data Curation, **IH**: Methodology, Software, **LN**: Methodology, Investigation, **KT**: Methodology, Investigation, **JM:** Methodology, Project administration**, RJM**: Methodology, Investigation, **KG**: Conceptualization, Writing - Review & Editing, Supervision, Project administration, Funding acquisition

## METHODS

### Animal care and surgery

#### Lesion

Following task training, Monkeys H and N underwent a stroke-induction and array implantation surgery. Preoperatively, animals were sedated with ketamine hydrocholoride (10 mg/kg), administered atropine sulfate (0.05 mg/kg), prepared and intubated. They were then placed on a mechanical ventilator and maintained on isoflurane inhalation (1.2-1.5%). Animals were positioned in a stereotactic frame (David Kopf Instruments, Tujunga, CA). A skin incision, bone flap, and dural flap were made over the lateral frontoparietal convexity of the hemisphere and the caudal region of the frontal lobe and rostral region of the parietal lobe was exposed unilaterally. After cortical exposure, the lesion was induced using surface vessel coagulation/occlusion followed by subpial aspiration ^22^. The lesion target was the forelimb region of primary motor cortex (M1) using anatomical landmarks.

Specifically, the lesion extended dorsally to a horizontal level including the precentral dimple (the lateral-most part of the of the M1 leg area) and ventrally to the central sulcus genu (the dorsal most part of the M1 face area). The central sulcus was slightly expanded, and the rostral bank of precentral gyrus was targeted constituting a lesion in the forelimb region of the “new M1” ^63^.

#### Microwire electrode array implantation in perilesional motor cortex

An 8x8 tungsten microwire multielectrode array with 500um electrode spacing and 375 um row spacing (Tucker-Davis Technology, Alachua, FL) was mounted to a micromanipulator, attached to the stereotax, and inserted into ventral premotor cortex (PMv) using anatomical landmarks (array placed just posterior to the inferior arcuate sulcus and lateral to the spur of the arcuate sulcus). Array reference and ground wires were tied to titanium skull screws that were placed anterior and posterior to the craniotomy respectively. Intraoperative recordings were monitored to determine depth of implantation (Monkey H: 1.8mm, Monkey N: 1.7mm).

#### Subcortical electrode array implantation

A vector array consisting of a stainless steel support body (419 um outer diameter) with a thin silicon shank (50uM thickness) with 32 electrode contacts (NeuroNexus, Ann Arbor, Michigan) was used to target subcortical sites. Specifically, a V1x32-Edge-15mm-100-177-CVZ32 with a 24mm support body, a 10cm flex cable, and 32 electrode contacts with 100 uM spacing spanning the last 3.1 mm of the silicon shank was used in Monkey H. The array was identical for Monkey N except the connector was different (Omnetics instead of ZIF clip) and the support body was 18mm instead of 24 mm.

For Monkey H, stereotaxic coordinates were used to plan the electrode trajectory to target the VL nucleus of motor thalamus. Before surgery, the site of insertion for the electrode trajectory was identified on the MRI and was measured with respect to visually identifiable landmarks such as the precentral dimple and central sulcus. During surgery, the electrode array was placed in a micromanipulator using a 3D printed adaptor designed to hold the array straight up and down and attached to the stereotax. The electrode was moved to stereotaxic coordinates. This insertion location was then verified by measuring its distances from the visually identifiable landmarks and confirming that they were as expected based on the pre-surgical measurements. The electrode was then lowered 24mm and secured in place.

Based on the imperfect insertion of the electrode array into motor thalamus in Monkey H, we used a different technique for the insertion of the array for Monkey N. Specifically, we adapted the following fiducial approach^64^ to ensure excellent alignment between surgical stereotaxic coordinates and MRI coordinates for more accurate targeting of the subcortical nuclei. We almost completely followed the approach outlined in Bentley et. al.^64^ with the following changes: Instead of using bone screws as fiducial markers, we used multi-modal fiducial markers (IZI Medical, Owings Mills, MD) that were placed on the skin during the subject’s MRI. Since these markers have a hole in the middle of them, skin tattoos were placed at the center of the fiducials for later measurement. In surgery, we measured the tattoos in stereotaxic coordinates. These measurements were used to align MRI coordinates to the surgical space’s stereotaxic coordinates. Surface landmarks that were pre-identified on the MRI (such as start of arcuate sulcus spur, anterior and posterior most aspects of the precentral dimple, anterior most point of the central sulcus) were measured in stereotaxic space to validate the coordinate transform. Our code and methodology for this realignment approach is described further https://github.com/smg211/NHP_Imaging.

An electrode trajectory that would reach the VL thalamic nucleus was then planned by converting the pre-identified MRI target coordinate to the surgical space stereotaxic coordinate. An angled trajectory was planned to increase the probability of hitting the target even considering errors of 1-2 mm. The surface entry location of the electrode and the electrode entry angle was selected. Electrophysiological recordings were monitored during the lowering of the probe, and spiking activity was observed while the electrode contacts were in cortex, midway through the lowering of the trajectory (likely recordings from caudate nucleus), and at the end of the trajectory.

Array reference and ground wires were tied to titanium skull screws that were placed anterior and posterior to the craniotomy respectively.

Following surgery, animals were administered analgesics and antibiotics, and were carefully monitored post-operatively for 7 days.

#### Microwire electrode array implantation in primary motor cortex

In neurotypical animals (Fig. 7, Monkeys B and F) an 8x8 tungsten microwire multielectrode array with 500um electrode spacing and 375 um row spacing (Tucker-Davis Technology, Alachua, FL) was mounted to a micromanipulator, attached to the stereotax, and inserted into right primary motor cortex (M1) using anatomical landmarks. Electrodes were lowered to a depth of 2 mm to target layer 2/3. Array grounds were tied to a titanium skull screw posterior to the craniotomy site, and array references were tied to titanium skull screw anterior to the craniotomy site. Intraoperative recordings were monitored to determine depth of implantation.

### Reach-to-grasp Behavior

#### Task

Prior to surgery, macaques were trained to perform a reach-to-grasp task that allowed us to automatically assess grasping behaviors of different difficulties throughout the course of behavioral recovery. Briefly the behavior required animals to reach out to grasp an object, and lift it to a certain height for a designated hold period in order to receive a juice reward. The animals had to fit their fingers through a “slot” to grasp the object, and the slot could be one of four shapes (Fig 1B). The shape of the slot changed the difficulty of the grasping task. This behavior is an adaptation of our previous “pinch-and-lift” task^21^.

As shown in Fig. 1A, animals initiate a trial by depressing a button for a designated hold time (Monkey H: 0.2 sec, Monkey N: 1.5 sec). Upon successful hold, a door blocking access to the slot and the object retracts (Supp. Fig. 2B-C), signaling the start of the trial. Animals then reached toward the task’s object. To successfully grasp the object, they had to shape their fingers to fit through a particular slot. On every trial, the slot could either be a “precision” slot, a “pinch” slot, a “tripod” slot, or a “power” slot (Fig. 1B). Each of these slot types constrained the grasp in different ways, making grasping easy (e.g. power slot in which a whole-handed grasp could be used to grab the object), more moderate (e.g. pinch or tripod slots in which only the index and thumb or index, middle finger, and thumb could fit into the slot to grasp the object), or very difficult (e.g. precision slot in which the index and thumb could fit into the slot in a particular configuration and in which the object had to be lifted out of the slot at a very particular angle). The slots were cut into a piece of acrylic that was shaped like a wheel, and were controlled by a motor that rotated the wheel such that the slot for that trial was aligned with the object (Supp. Fig. 2D-E). The wheel was rotated into the correct position prior to the start of each trial. Finally, animals lifted a 3D printed object that was a tall rectangle. The object was 12.5mm in width, 7mm in thickness, and 45mm in height. The object needed to be lifted approximately 15mm to trigger the completion of an infrared (IR) beam break sensor (3mm LED, Adafruit.com). Animals had to hold the device at the lifted height for 0.267 sec, following which they received a juice reward through a chair-mounted juice tube. Animals had 10 seconds from the time of the door opening to complete the object lift. If the object was lifted high enough to allow the IR beam sensor to complete, but did not maintain that hold for long enough (i.e, “hold error”), the 10 second timer was restarted. If animals did not complete the lift by the timeout time, the trial would end and the door would gently close. Animals performed 10-100 trials per day, depending on impairment. Which slot types were used on which day and their relative ratios of presentation were selected based estimates of impairment.

Two cameras (CM3-U3-13Y3C-CS, Point Grey, Richmond, BC, Canada) were mounted to a stainless steel platform providing a side and top view of the animal performing the reach-to-grasp task. Camera frames were captured at 50 Hz. Camera frames and task sensor activity were synchronized using the electrophysiology recordings system (Tucker-Davis Technology, Alachua FL).

#### Automatic extraction of behavioral timepoints

Trials were automatically scored as “rewarded” or “unrewarded” based on whether the object was lifted above the 15mm height threshold for the 0.267 second hold period.

Behavioral markers were annotated from a combination of the video data and task sensor data.

*Reach start* and *grasp start* were determined based on kinematics that were extracted from the side-view camera data with DeepLabCut ^65^ using the following approach:

1. A DeepLabCut network was trained to identify the metacarpophalangeal (MCP) joint of the index finger in a subset of frames. The network error on training data was 2.04 pixels (2.76 pixels) and on test data was 2.35 pixels (5.65 pixels) for Monkey H (Monkey N) .
2. For each trial, animal hand position was tracked using the metacarpophalangeal (MCP) joint of the index finger. On frames where the likelihood of identifying the index MCP position was less than 0.9, the position was marked as “NaN”.
3. The positions of the index MCP between the time of the door opening and end of the trial were collated. The x and y positions were temporally smoothed using a gaussian kernel of length 210ms (11 frames) with a standard deviation of 20ms (1 frame). In code: window = scipy.signal.gaussian(11., 1.0). NaN values were ignored in the temporal smoothing.
4. The hand speed was calculated from the smoothed index MCP trajectory.
5. The frame in which the index MCP crossed the line defined by the front panel of the task apparatus (black dashed line in Supp. Fig. 2G) was identified as “*crossing frame*”. This crossing corresponded to a point mid-way through the animals’ reach.
6. The peak hand speed in a [-200ms, 200ms] window centered around the *crossing frame* was identified as the “*peak hand speed”* and the time of the *peak hand speed* was identified as *peak hand speed frame*.
7. The hand speed trajectory was traversed backwards in time from the *peak hand speed frame* to identify the frame at which the hand speed was less than 0.1 × *peak hand speed*. The frame was marked as “*reach start frame”*.
8. Similarly, the hand speed trajectory was traversed forwards in time from the *peak hand speed frame* to identify the frame at which the hand speed was less than 0.1 × *peak hand speed*. The frame was marked as “*grasp start frame”*.
9. *Reach start frame* and *grasp start frame* were converted to *reach start* and *grasp start* times in seconds using the synchronized pulses in the neural recording system that coincided with each captured camera frame.

Trials in which there was no *crossing frame* identified were not analyzed for *reach start* or *grasp start* times. All trials were manually reviewed after this procedure to ensure proper identification of *reach start* and *grasp start* times.

The DeepLabCut network was also designed to track the movement of the sliding door. The time at which the door retracts at the start the trial was marked as *door open frame* and was converted to *door open* times in seconds.

The time at which the IR beam break sensor first showed that the object was lifted above the 15mm height criteria was designated as “*first beam lift”*. The time at which the object was lifted above the 15mm height criteria for the last time in the trial (typically before the reward is delivered) was designated as “*last beam lift”*.

The time at which the IR beam break sensor was lifted above the 15mm height criteria for 0.267 seconds was designated as “*reward time”.* We note that due to slight imprecision in the task code apparatus (each task cycle was not exactly the same length of time), some trials that were designated as reward trials by post-processing analysis of the beam break sensor were not actually rewarded during the behavior, and some trials that were rewarded during the behavior were not designated as reward trials by post-processing analysis. For consistency, we use the post-processing analysis of the beam break sensor to assign trials to as rewarded versus unrewarded.

#### Calculation of behavioral metrics

Reach duration is defined as the time between *reach start* and *grasp start*. Grasp duration is defined for rewarded trials only, and is the time between *grasp start* and *reward time*. Normalized reach duration and grasp duration is used in order to combine reach and grasp durations across trials types of different difficulty. Normalized reach duration or normalized grasp duration for a particular trial is defined as reach or grasp duration divided by the average reach or grasp duration of all the pre-stroke trials that were executed to that trial’s particular slot. For example, the normalized reach duration of a trial performed with the “pinch” slot would be calculated as the reach duration of the trial divided by the mean reach duration for all pre-stroke trials completed on the pinch slot. When normalized times are > 1, this indicates reach or grasp durations that are much slower than pre-stroke behavior. When normalized times ∼= 1, this indicates reach or grasp duration that are similar to the pre-stroke behavior. In Fig. 1E, F, only rewarded trials are used to calculate normalized reach duration and normalized. grasp duration.

Normalized trial times are also calculated using the time between *reach start* and *last lift* and normalizing using pre-stroke baseline data as outlined above (Fig. 4EF)

Percent correct is calculated as the fraction of rewarded trials for a particular day.

To combine the normalized reach and grasp durations and the percent correct metrics, a “recovery index” metric was developed. The recovery index was defined by first calculating the total “trial time” for each trial. Trial time for rewarded trials was defined as the time between the reach start and reward time. Trial time for unrewarded trials was defined as 10 seconds (the timeout time). Then, the normalized trial time was calculated by dividing each trial’s trial time by the average pre-stroke trial time for that trial’s slot. The normalized trial times were then averaged over all trials on each day to yield an average normalized trial time for that day. Note that normalized trial times will typically vary between 1 (indicating similar performance to pre-stroke) and a larger number (indicating longer durations compared to pre-stroke). The day-averaged normalized trials times were then all decremented by 1 (such that now values varied between zero and a larger number), and then divided by the first post-stroke day’s value (such that now values varied between 0 and 1 where 0 indicated pre-stroke baseline, and 1 indicated deficits as large as the first day of behavioral testing after stroke). Finally, the values were subtracted from 1, such that a value of 0 indicated a deficit as large as the first day post-stroke, and a value of 1 indicated behavior similar to the pre-stroke baseline. The average recovery index over all trials on a particular session is used as that day’s recovery index. Recovery indices are plotted in Fig. 1G.

In some analyses involving spiking activity, recovery curves are split into “early” and “late” segments. For monkey N, “early” sessions are designated as sessions where the recovery index is < 0.5, and “late” sessions are designated as sessions where the recovery index is > 0.5. For monkey H, only two sessions (and only one with useable neural data) would have fallen in the “early” category if a threshold of 0.5 was used. Thus, the threshold for “early” and “late” was set to 0.8. See supplementary table 1 for which days are early vs. late.

#### Definition of reaching and grasping epochs

The reaching epoch is defined as the time between *reach start* and *grasp start*. Only trials in which *reach start and grasp start* were both defined were used for reaching epoch analyses. The grasping epoch is defined as the time between *grasp start* and *reward start* if a trial is rewarded, or *grasp start* and *(door open + 10 sec)*, i.e. the trial timeout time, for unrewarded trials.

### Sequential target-reaching task

The sequential target-reaching task (Fig. 7a) was implemented on a 15-inch touchscreen tablet (Galaxy Tab S8 Ultra, Samsung). Touchscreen input was sampled at 120 Hz and recorded on the task workstation. At the start of each trial, the screen displayed 9 yellow unfilled circles (diameter = 3.8 cm), arranged in a symmetrical 3 × 3 grid (inter-row and inter-column spacing = 6.5 cm). To initiate a trial, animals held their hand on an external start button positioned below the screen for 1 s. Then, one of the yellow circles on the screen would become filled, representing the first target. Immediately after the animal touched the target, that target would become unfilled and the next target would be filled. This process repeated until the animal touched the fifth target. A tone was played when each target was presented. Immediately after the animal touched the fifth target, a separate tone was played, the screen color changed to all white, and a juice reward was automatically dispensed. After juice was dispensed, the screen color changed to all black, and the inter-trial interval (ITI) started. The ITI duration was randomly selected from a normal distribution with *μ* = 1.0 s and *σ* = 0.2 s. After the ITI, a start tone was played indicating the start of the button-hold period of the next trial.

#### Extraction of wrist kinematics

Two cameras (CM3-U3-13Y3C-CS, Point Grey, Richmond, BC, Canada) were mounted to a stainless steel platform providing a side and top view of the animal performing the sequential target-reaching task. Camera frames were captured at 50 Hz. Camera frames were synchronized using the electrophysiology recordings system (Tucker-Davis Technology, Alachua FL). Wrist position was tracked on the side camera by training a Deeplabcut network, which was then used to compute wrist speed.

#### Definition of reaching epoch for Monkeys B, F

The reaching epoch is defined as the time between [-1 sec + *first_reach start*] to the *target_touch* event of the last (fifth) reach. *First_reach_start* was determined based on a beam breaking sensor that was embedded in the external start button. When animals had the button depressed, the beam break sensor was triggered. When they began their first reach the beam break sensor became untriggered. *First_reach_start* was defined as the transition from the triggered to untriggered state. *Target_touch* was defined as the time at which the touchscreen registered a touch inside the presented target.

### Neural signal processing and analysis

#### Local field potential (LFP) processing

LFP data was acquired from both the perilesional motor cortex and subcortical electrode arrays at 3051.8 Hz using Analog PZ5 Pre-amplifier, and the RZ2 Bioamplifier (Tucker-Davis Technology, Alachua, FL). Offline, signals were downsampled by a factor of 3 using scipy.signal.decimate (which downsamples after applying an anti-aliasing filter). For stroke animal data, trial segments were extracted in windows of [-3, 10] seconds with respect to reach start time and for neurologically intact animal data, trial segments were extracted in windows of [-1, 5] seconds with respect to first reach start time. On each day, trials were manually inspected alongside camera data for any movement or chewing artifacts (note that animals were not head-posted) and were discarded from neural data analysis if there were any artifacts. Using the remaining trials, all channels were manually inspected and discarded if there were unusually large, intermittent fluctuations in signal. Note that LFP data was *not* referenced beyond hardware referencing to the reference and ground skull screws.

#### Power spectral densities

Power spectrums of trials and channels that were not discarded were calculated using scipy.signal.spectrogram in windows of 4 seconds with a window overlap of 1 second. Spectrograms were averaged over channels and time and plotted (Fig. 1H-I).

#### Beta window selection

The beta band window was selected to be a ∼20 hz width window centered on the “beta bump”. Since animals showed slightly different beta band frequencies, the beta band range was shifted to be animal-specific. The selected ranges of 15-35 Hz (Monkey H), 10-30 Hz (Monkey N), 18-30 Hz (Monkeys B and F) were also validated using an automated method for parameterizing power spectra into periodic and aperiodic components^23^. For each day, the beta band was identified using the channel-averaged and trial-averaged power spectra. Across all days, the limits of identified beta band spectra was within and spanned the selected beta ranges for each monkey.

#### Beta burst identification and burst feature calculation

In order to identify beta bursts the following procedure was followed:

1. First the channel data was bandpass filtered in the animal-specific beta range (15-35 Hz for Monkey H, 10-30 Hz for Monkey N, 18-30 Hz for Monkeys B and F) using a 3^rd^ (4^th^) order Butterworth bandpass filter for Monkeys H and N (B, F).
2. Then the amplitude of the filtered signal was calculated using the Hilbert transform (scipy.signal.hilbert)
3. The amplitude of the complex valued signal was calculated, *amp_ch_day* (np.abs)
4. For each channel on each day, three parameters were calculated:

a. *Sigma_ch_day*: the standard deviation of *amp_ch_day*
b. *Thresh_ch_day*: the median of *amp_ch_day*
c. *Thresh2_ch_day*: the median *amp_ch_day* plus *sigma_ch_day*
5. Burst events on each channel were detected by scanning through *amp_ch_day* and identifying epochs where the following two criteria were satisfied:

a. *amp_ch_day* exceeded *Thresh_ch_day* for at least 100 ms
b. the maximum of *amp_ch_day* exceeded *Thresh2_ch_day*

At least three cortical channels must be bursting at a given timepoint to classify the timepoint as part of a “cortical burst event”.

For each burst event, the following features were calculated:

1. Normalized amplitude – the average amplitude during the burst event minus Thresh_ch_day and divided by Sigma_ch_day
2. Duration – length of burst event
3. Frequency – the average of the instantaneous frequency of the burst event. Instantaneous frequency *inst_freq_ch_day* was calculated by computing:

a. the differentiated unwrapped phase of the complex valued signal (after the Hilbert transform using np.angle, np.unwrap, np.diff), yielding *dphase_ch_day*
b. normalizing the phase: *inst_freq_ch_day* = *dphase_ch_day* * (1 / 0.5*pi) * *fs*;

#### Definition of frac_thresh_global_local

To identify the threshold of fraction of cortical channels bursting used to distinguish local and global events, the following procedure was followed:

1. For all timepoints and trials on a given day, the fraction of cortical channels that are bursting is calculated. All timepoints that are part of cortical bursts events are extracted.
2. The distribution of cortical channels bursting is calculated (as shown in Fig. 2A).
3. The average of the distribution across all days analyzed is calculated (i.e. average of all colored lines in Fig. 2A)
4. A gaussian mixture model with two Gaussians is fit to the average distribution
5. The intersection point between the two identified distributions is marked as the *frac_thresh_global_local* for all days.

#### Identification of local and global burst events

At each timepoint, the fraction of cortical channels that are bursting is calculated. If the fraction exceeds *frac_thresh_global_local*, the timepoint is marked with a “2” for global. If the fraction is less than *frac_thresh_global_local* and the number of channels bursting is greater than or equal to three channels, the timepoint is marked with a “1” for local. Else, the timepoint is marked with a “0” for non-bursting. The timeseries of 0s, 1s, and 2s is then processed. Local burst events are defined as continuous epochs of 1s that are at least 100ms in duration. Global burst events are defined as continues epochs of 2s that are at least 100ms in duration.

#### Single unit extraction and processing

Raw neural signals were acquired at 24414.0625 Hz using Analog PZ5 Pre-amplifier, RZ2 Bioamplifier, and RS4 Data Streamer (Tucker-Davis Technology, Alachua, FL). Offline, signals were median subtracted (median computed on each bank of 16 channels in array) to reduce motion artifacts and external noise and then bandpass filtered between 300 Hz and 6000 Hz. Spikes were then extracted from signals using MountainSort software^66^. Signal-to-noise (SNR) ratios of the extracted unit waveforms were re-calculated after sorting. SNR is defined as the peak minus trough of the average waveform divided by the standard deviation of the waveform (excluding the region between slightly before the trough and slightly after the peak). Artifactual units were manually tagged for removal after processing. Some units were combined after manual inspection to avoid duplication of the same unit multiple times in the analysis. After curation, all waveforms with SNR metrics > 3 and mean firing rates > 0.5 Hz across the entire recording were visually inspected and selected for analysis. Spike counts were binned in 10ms bins and converted to a firing rate.

#### Trial-averaged firing rate and firing rate variability during reaching and grasping epochs

To calculate the trial-averaged firing rate and trial-averaged firing variability during reaching, trials with *reach start* and *grasp start* defined were noted. Each unit’s mean firing rate between *reach start* and *grasp start* was calculated for each trial, and the trial-average firing rate was then computed across trials for a given session. Similarly for firing rate variability, each unit’s firing rate variation between *reach start* and *grasp start* was calculated for each trial, and the trial-average firing rate variation was then computed across trials for a given session. The same procedure was used for grasping, except that the mean rate and rate variation was calculated between *grasp start* and *reward start* if a trial is rewarded, or *grasp start* and *(door open + 10 sec)*, i.e. the trial timeout time, for unrewarded trials.

#### Entrainment to beta events

To calculate each units’ entrainment to local and global beta events during different parts of the task, the following procedure was followed:

1. Spiking activity was binned in 1ms bins
2. Local and global beta burst events were identified as detailed above in “*Identification of local and global burst events”*
3. For each LFP channel, the timeseries of beta phases was extracted using np.angle applied to the hilbert transformed data.
4. For entrainment calculated for a particular behavioral event (e.g. grasping entrainment), beta burst events that occurred during that behavioral event were selected.
5. Whenever a unit spiked during these beta burst events, the phase of the beta signal from each channel was collected.
6. After this process, each unit had an nSpikes_loc x nChannels data array for local beta bursts and an nSpikes_glob x nChannels data array for global beta bursts, where each entry corresponds to the phase of the LFP channel when that unit spiked. nSpikes_loc and nSpikes_glob indicate the number of times that unit spiked during local and global beta bursts that occurred within the particular behavioral event.
7. For each unit and each LFP channel, the strength of entrainment was calculated by calculating the resultant vector^67^ of the phases, yielding an nChannels x 1 vector for both local and global beta bursts for each unit.
8. To test the significance of each of these unit-channel entrainment strengths, the following procedure was followed to build a null distribution of expected entrainment given the number of spikes, and the phase distribution of the LFP:

a. For all timepoints where there was a beta burst event that fell within the behavioral event, the phases of the LFP channel were collected.
b. The number of spikes that fell within a beta burst event and within the behavior event were counted, yielding nSpikes_loc or nSpikes_glob depending on whether local or global bursts were being analyzed.
c. A shuffle distribution of resultant vector lengths was then built. For each shuffle, the nSpikes samples from the LFP channel phase distribution built in (a) were drawn, and the resultant vector length was calculated.
d. This procedure was repeated 100 times, yielding a distribution of expected resultant vector lengths given the firing rate of the unit during beta bursts of the behavior.
9. Individual unit-channel resultant vector lengths were considered “significant” if they exceeded the 95^th^ percentile of the shuffle distribution for that unit-channel pair.
10. “modulation z-score” values that report entrainment strength were calculated as the true resultant vector length of the unit-channel pair minus the mean and divided by the standard deviation of the shuffle distribution.

#### Correlation between entrainment to beta events and change in firing rate and variation during beta events

For each unit, significantly entrained unit-channel pairs were identified and the median of the modulation z-score was computed across these channels. This median modulation z-score was paired with the difference between the trial-averaged firing rate and trial-averaged firing rate variability during grasping that was computed during beta burst events and non-beta burst events.

#### Population dynamics model

To quantify how global and local beta oscillations influence “temporal predictability” of neural population dynamics, a linear dynamics model was employed. Neural population activity at the next time point (*X_t_*) was modeled as a linear function of current neural population activity (*X_t_* = *AX_t_*_-1_). The variance explained by this model indicated how well the model could estimate future fluctuations in neural activity based on current state of neural activity i.e, “temporal predictability”.

Models were fit on spiking data binned in 50ms bins, and included 0.5 seconds before reach start to the time of the first object lift. Only units that were significantly modulated during reach or grasp were included in the population activity. In order to control for the amount of data and number of detected units that varied across days, a minimum number of trials and units was identified for each animal. For Monkey H (N), each model was fit using 8 (9) rewarded trials and 12 (26) neural units. Trials and units were subselected randomly, and model fitting was repeated a total of 200 times. Trials that were not used in the fitting were held out and predicted by the fit model.

An example of model predictions that have been trial-averaged are shown in Figure 5G, *bottom* in comparison to the true data that was trial-averaged.

#### Comparisons of population dynamics model R^2^ during global, local, and no bursts during grasping

True spiking and model-predicted spiking data between grasp start and reward (or trial cue plus 10 seconds for unrewarded trials) were aggregated depending on whether they occurred during no bursts, global beta bursts, or local beta bursts. For each trial, all bins during the grasping epoch corresponding to “no bursts”, “global burst”, and “local burst” were used to calculate a trial-R2 metric for “no bursts”, “global bursts”, and “local bursts” respectively.

#### Spatial clustering of channels during local and global bursts

For each burst *i* that is detected during reaching and grasping phases of movement, the burst center is calculated and the average distance of all channels from the burst center is calculated. The burst center (*loc center_burst_*_=*i*_ ∈ *R*^2^) is calculated as a weighted mean of the (x, y) location of all channels that are active during the burst (*loc_burst_*_=*i, channel*=*j*_ ∈ *R*^3^) weighted by the fraction of time that channel is active during the burst (0 ≤ *frac act_burst_*_=*i,channel*=*j*_ ≤ 1):

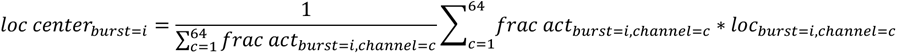

The average distance to the burst center (*dist center_burst_*_=*i,*_ ∈ *R*^1^) is calculated as a weighted mean of the distance of each channel to the burst center weighted by the fraction of time that channel is active during the burst (*frac act_burst_*_=*i,channel*=*j*_):

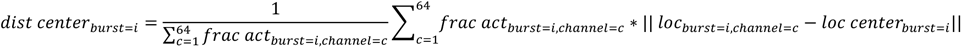

For each detected burst event, there is a single *loc center_burst_*_=*i*_ and *dist center_burst_*_=*i*_.

To detect if the distances to the burst centers are spatially distributed (Fig. 3A, *left*) or spatially clustered (Fig. 3A, *right)*, we computed what distances would be if channels composing each burst were shuffled in time so as to preserve the average number of channels active in a burst and the average activation of each channel across local bursts. Specifically, all binary arrays that indicated which channels were bursting (*burst*_i_ ∈ *nCh x nTime_i_*) were collated from either reaching or grasping epochs ([*burst*_1_, *burst*_2_, … *burst_i_*, … *burst_N_*] ∈ *nCh x* ∑^*N*^_*i*=1_ *nTime_i_*) where *N* is the number of bursts on a given session within the reach or grasp epoch. This large array was then shuffled along the columns dimension (i.e. along the time dimension), and then re-segmented into individual burst events. The *loc center_burst=i_* and *dist center_burst=i_* where then computed for each re-segmented, shuffled burst event.

The shuffling procedure was repeated 100 times, yielding a distribution of *dist center_burst=i_* for each session and set of bursts (i.e. reaching or grasping epoch bursts). The number of datapoints in the distribution is *N* ∗ 100.

Finally, to detect if the distances to the burst centers are spatially distributed (Fig. 6A, *left*) or spatially clustered (Fig. 3A, *right)*, the distribution of true distances to the burst center for a particular session and set of bursts (i.e. reaching or grasping epoch bursts) were centered by the mean of the shuffle distribution for that session and epoch, yielding the quantity “distances from burst center (mm) – shuffle mean” (Fig. 3C). We then tested whether this distribution of shuffle-subtracted distances was significantly lower than zero (Fig. 3C). Distributions of global and local spatial distances were compared directly (Fig. 3D).

---

**Supplemental Table 1:** Days used for Behavioral, LFP, and spiking analysis:

| Monkey H | Behavior<br>(rewarded<br>trials) | LFP | Spikes | Early<br>/late | Monkey N | Behavior<br>(rewarded<br>trials) | LFP | Spikes | Early<br>/late |
| --- | --- | --- | --- | --- | --- | --- | --- | --- | --- |
| Day 7 | X (x) | X | X | Early | Day 7 | X | X | X | Early |
| Day 8 | X (x) | * | * | Early | Day 8 | X (x) | X | X | Early |
| Day 10 | X (x) | X | X | Early | Day 10 | X (x) | X | X | Early |
| Day 11 | X (x) | X | X | Early | Day 11 | X (x) | ** | ** | Early |
| Day 13 | X (x) | X | X | Late | Day 14 | X (x) | X | X | Early |
| Day 14 | X (x) | X | X | Late | Day 15 | X (x) | X | X | Late |
| Day 16 | X (x) | X | X | Late | Day 17 | X (x) | X | X | Late |
| Day 20 | X (x) | X | X | Late | Day 18 | X (x) | X | X | Late |
| Day 21 | X (x) | X | X | Late | Day 73 | X (x) | X | **** | Late |
| Day 23 | X (x) | X | X | Late | Day 74 | X (x) | X | **** | Late |
| Day 24 | X (x) | X | X | Late | Day 80 | X (x) | *** | **** | Late |
|  |  |  |  |  | Day 81 | X (x) | *** | **** | Late |
|  |  |  |  |  | Day 85 | X (x) | *** | **** | Late |
| **Total**<br>**sessions:** | **11** | **10** | **10** |  |  | **13** | **9** | **7** |  |

\* Significant head movements that eventually broke the subcortical array connector and impaired spiking detection. Connector was repaired on Day 9.
\*\* No neural data recorded during this day due to animal movements preventing headstage clipping
\*\*\* Subcortical array connector broken
\*\*\*\* No spiking activity detectable on cortical array during online behavior sessions

**Supplemental Figure 1:**
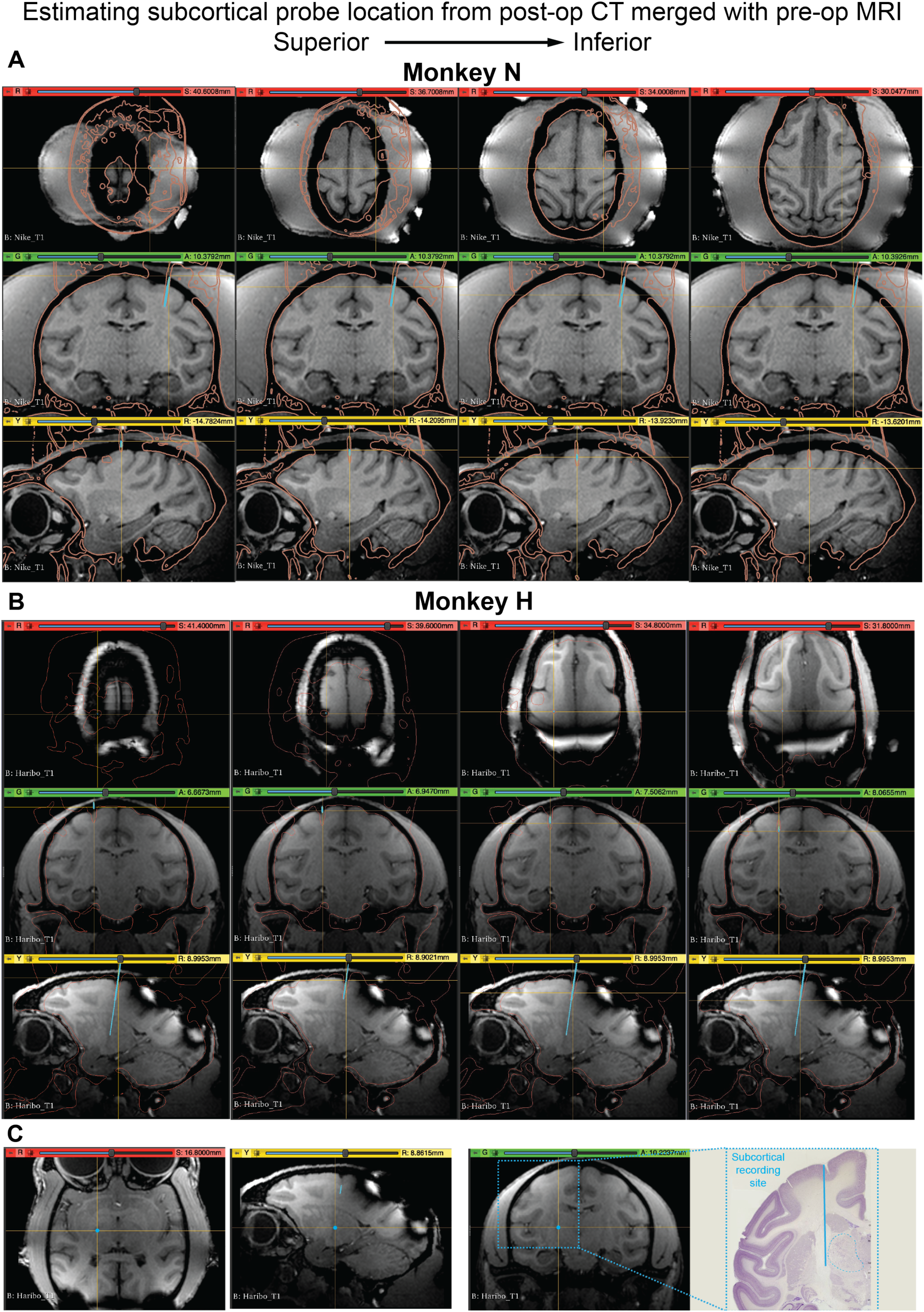
Estimating the subcortical probe location. Prior to surgery, cranial 3T MRI images were acquired for both animals using a T1-weighted sequence. After surgery, cranial computed tomography (CT) images were acquired for both animals. The MRI images were aligned and resliced in ACPC coordinates using FieldTrip . Then, both the CT and re-sliced MRI images were loaded into 3D Slicer. The CT images were windowed using the default CT-bone window and were thresholded to extract a volume containing the skull. This volume also contained the implanted microelectrode array in perilesional cortex and support body of the subcortical electrode array. **A, B:** The extracted skull and implanted electrode volume (CT, orange) was then manually aligned with the MRI such that the CT skull aligned maximally with the skull in the MRI image. When traversing the axial axis of the scan, the CT skull volume shows excellent alignment to the MRI in the axial, coronal, and sagittal axes in places where the skull was not disrupted due to the surgical intervention. After alignment, a 3D model of the subcortical electrode array was imported into the 3D slicer image (cyan). The array was manually aligned such that the support body of the array maximally overlapped with the part of the CT volume corresponding to the support body. The silicon shank part of the subcortical electrode array was not visible on the CT. The alignment between the support body and the CT volume was verified by scanning through the axial, sagittal, and coronal axes and can be seen in **A, B**. The final location of the recording electrodes was then visualized in **C** (Monkey H), and **Fig 1D** (Monkey N). T1-weighted sequences do not visualize subcortical nuclei particularly well. To estimate the structures where the subcortical electrodes were positioned, brain slices from the Brainmaps.org atlas were visualized^69^. Slices that best matched the cortical anatomy of the coronal MRI slice where the electrodes were placed (shown in C and Fig. 1D) were selected (Monkey N: Atlas slice 8.2mm AP, Monkey H: Atlas slice 10.0mm AP). These slices were within 1-2mm of the AP axis of the animals’ MRI (Monkey N: 9.9mm AP, Monkey H: 10.2mm AP). The electrode trajectories were estimated on the atlas slices by computing the ML and DV coordinates of the final electrode position and the angle of the electrode tract in the MRI. The angle and the final electrode position dictated the electrode trajectory shown in Fig. 1D and **C**. These slices reveal that the final electrode positions were either in (Monkey N) or just outside in the white matter (Monkey H), the VL nucleus of the motor thalamus. For the purposes of this study, we simply refer to these electrode recording locations as “subcortical”.

**Supplemental Figure 2:**
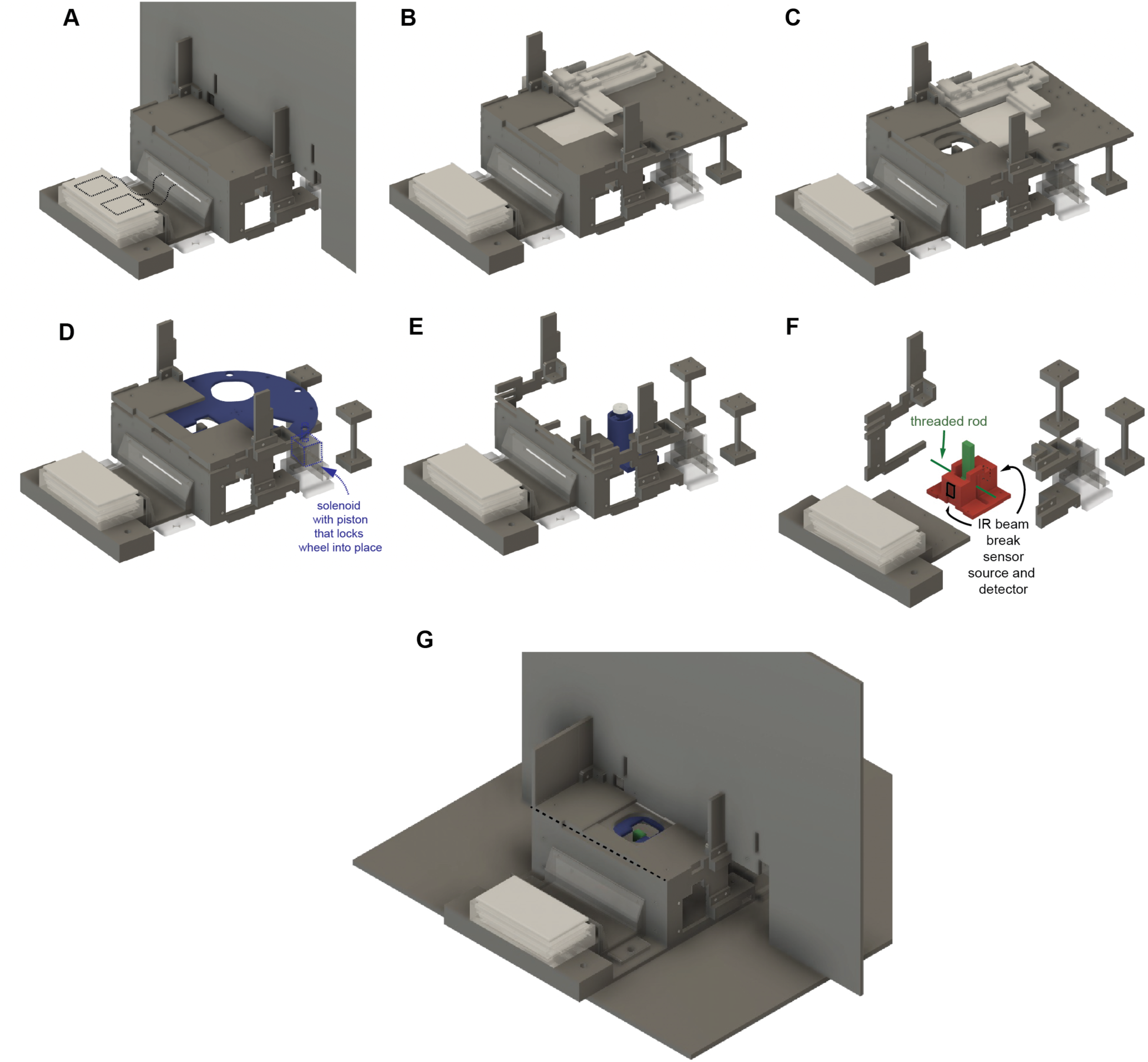
Reach-to-grasp task The CAD model of the reach-to-grasp task. A) Appearance of the fully assembled reach-to-grasp task from the perspective of the side camera at the start of a trial. The button is highlighted in white. The button houses two square force sensitive resistors (Sparkfun.com, SEN-09376) with wires that are routed to the back of the task (indicated by dashed black lines). The wires are covered by a “wire protector” piece. Animals must first depress the button for a designated hold period in order to open the door to the slot and object to start the trial. B) Removal of “backsplash” for better visualization of the slot door apparatus (shown in white). A linear slide potentiometer (Bourns PSM60-081A-103B2) is seated behind the backsplash and has a lever that is fixed to a 3D printed door. C) After the button has been depressed for the designated hold period, the door slides open, revealing the object and the slot that must be reached through in order to grasp the object. D) Removal of the door and the structural pieces of the task apparatus for better visualization of the “wheel” that has the slot cutouts. The wheel is fixed to a DC motor through a coupler (shown in E). Prior to each trial, the motor spins the wheel such that the correct slot is aligned with the object. A magnetic solenoid’s (Adafuit.com, Product ID = 3992) piston is released through holes on the outside of the wheel to lock the wheel in place and prevent the animal from moving the wheel during a trial. The tripod slot is shown in the current trial. E) The wheel and other structural pieces of the task have been removed to visualize the DC motor (Adafuit.com, Product ID 4416), the DC motor coupler (Pololu Robotics and Electronics, Pololu item # 1079, Pololu Universal Aluminum Mounting Hub for 3mm Shaft, #2-56 Holes (2-Pack), and the housing that was 3D printed to attach the motor to the task apparatus. F) Removal of the wheel motor and other structural parts to reveal the “object” (green), and the custom 3D printed apparatus that houses the object (red) and the IR beam break sensors that detect the object height. The object has a threaded rod that twists through its body, and through vertical slots on the housing. This rod and the track for the rod in the housing constrains the motion of the object to be up and down. The housing also contains cutouts for the IR beam break sensors (Adafruit.com, Product ID 2167) that detect if the object has been lifted above the 15mm height (shown in black). G) View of the re-assembled task with the door open revealing the slot and the object to be lifted. The black horizontal line indicates a line used in behavioral analysis (Methods, Reach-to-grasp behavior, automatic extraction of behavioral timepoints, relevant to identification of the “crossing frame”)

**Figure S3:**
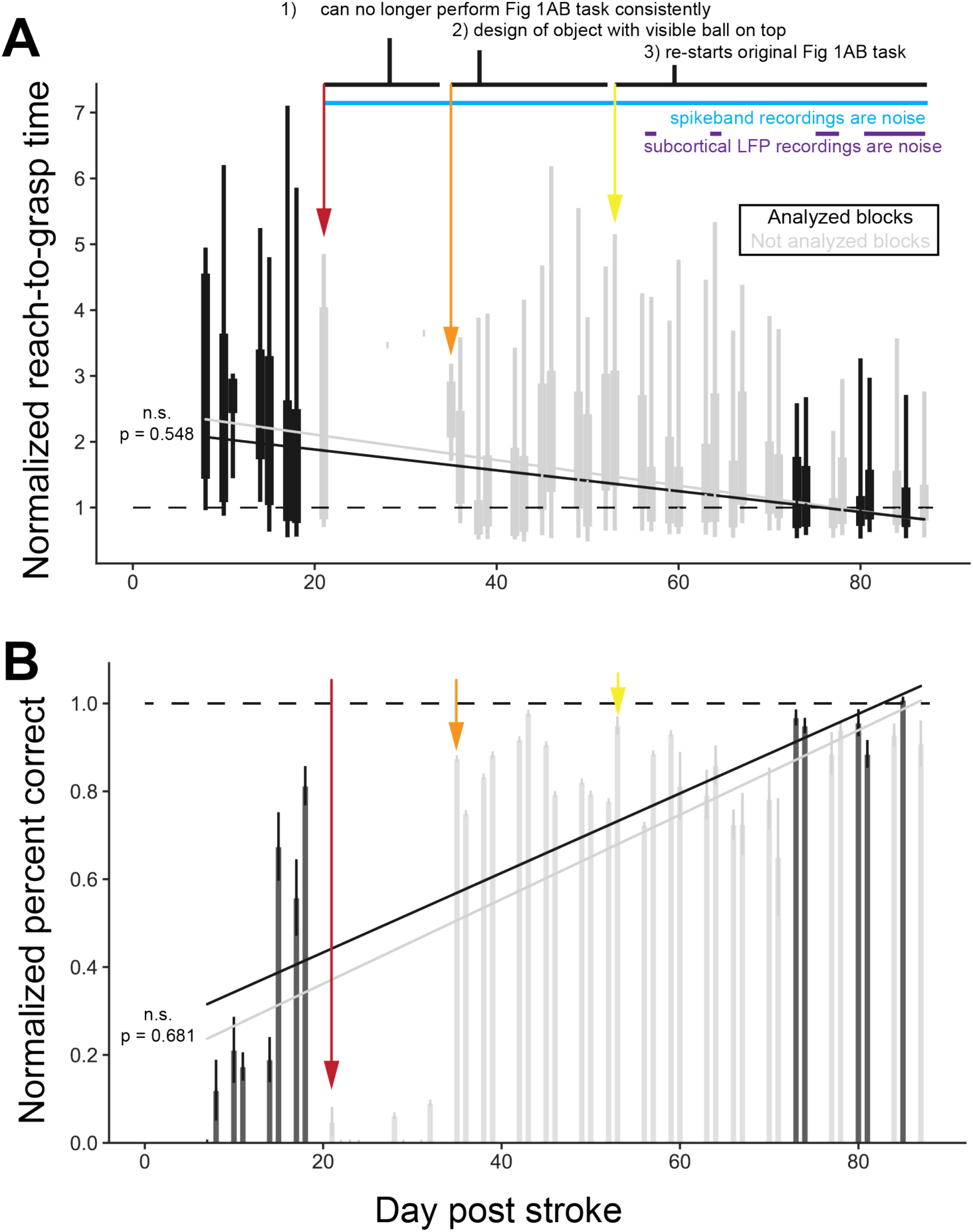
Monkey N full recovery curve. Monkey N underwent an unusual recovery curve. Initially, from days 7 to 18 after stroke, Monkey N exhibited a typical recovery with a reduction in reach-to-grasp time and an increase in percent correct. At Day 21 (red arrow), we noticed a return of his arm and hand impairment concomitantly with a significant reduction in cortical array spiking quality. We performed neuroimaging (CT) but were unable to see evidence of an obvious secondary brain injury or of an infection. We continued attempting to test Monkey N using the reach-to-grasp task outlined in Fig 1AB, but he continued to exhibit an inability to perform the task and poor cortical array spiking signals. At Day 35 (orange arrow) we introduced a new “object” within the same reach-to-grasp task format. The new object had a large ball on top of the existing rectangular object, which made finding the object and grasping it much easier. We continued testing Monkey N on this task until Day 53 at which point he was able to begin performing the original reach-to-grasp task again. We are still investigating the possible causes of his secondary decline. Because of this unusual recovery curve and because we have been unable to pinpoint the cause of the secondary decline, we have chosen to focus on the initial albeit incomplete recovery curve (days 7- 20) augmented by a few days at the end of his recovery once his performance was back at pre-stroke baseline. Since the purpose of our study is to examine how global and local beta dynamics change with recovery from stroke, it was important to us that our choice of sessions did not modify the overall dynamics of Monkey N’s recovery curve. To test this, we fit a linear regression to behavioral data from all days (gray lines in A, B) and fit a regression to only the sessions we analyzed in Fig 1EFG (black lines in A, B). We tested whether the slopes of these regressions differed significantly (F-test). For both normalized reach-to-grasp time and normalized percent correct there was no significant difference between the slopes of the gray and black lines (A: normalized reach-to-grasp time, p = 0.548, B: normalized percent correct, p = 0.681). See Table S1 for more details about exactly which days of data were used in each analysis.

**Figure S4:**
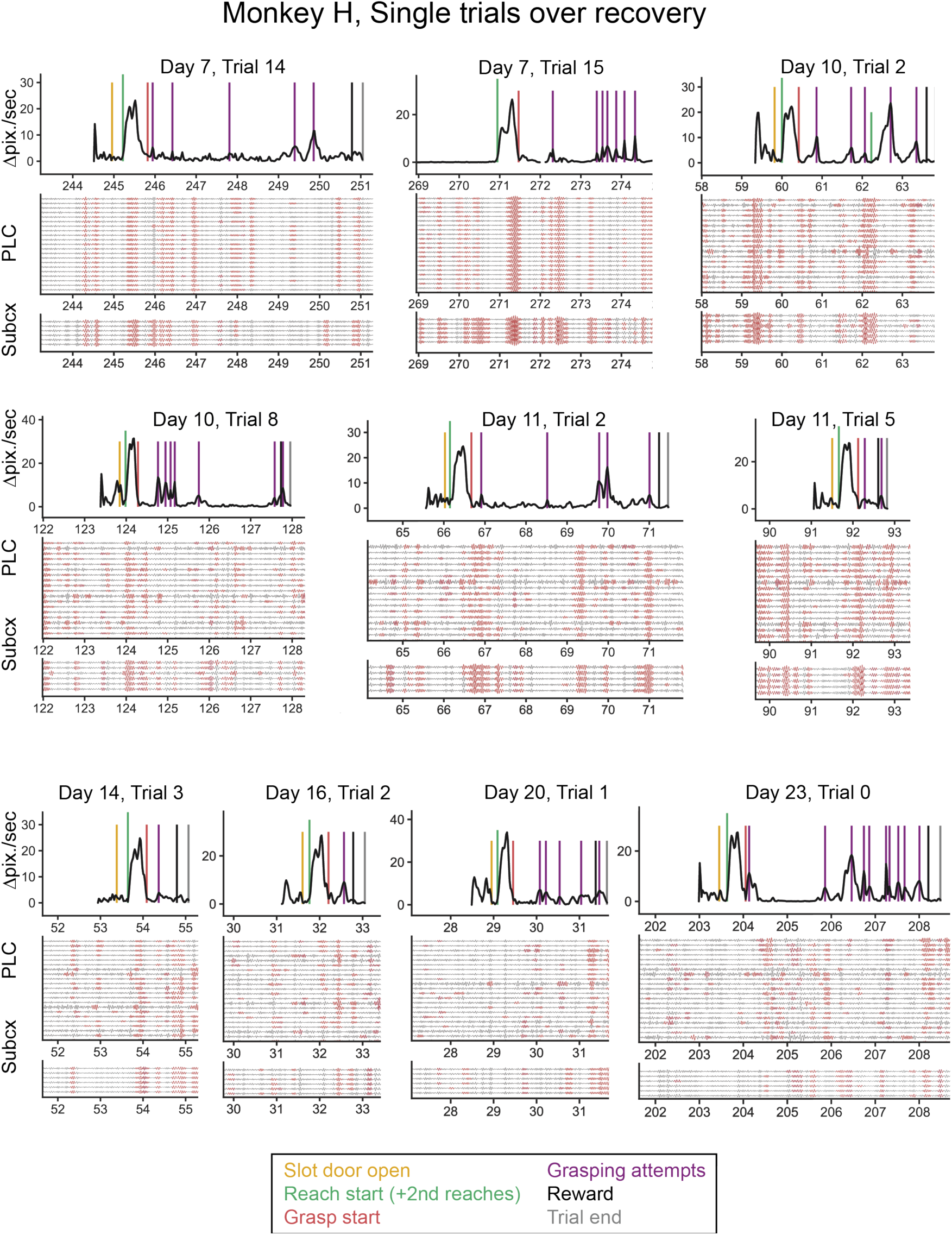
Monkey H single trials over the course of recovery. In the same style as Fig. 1JK.

**Figure S5:**
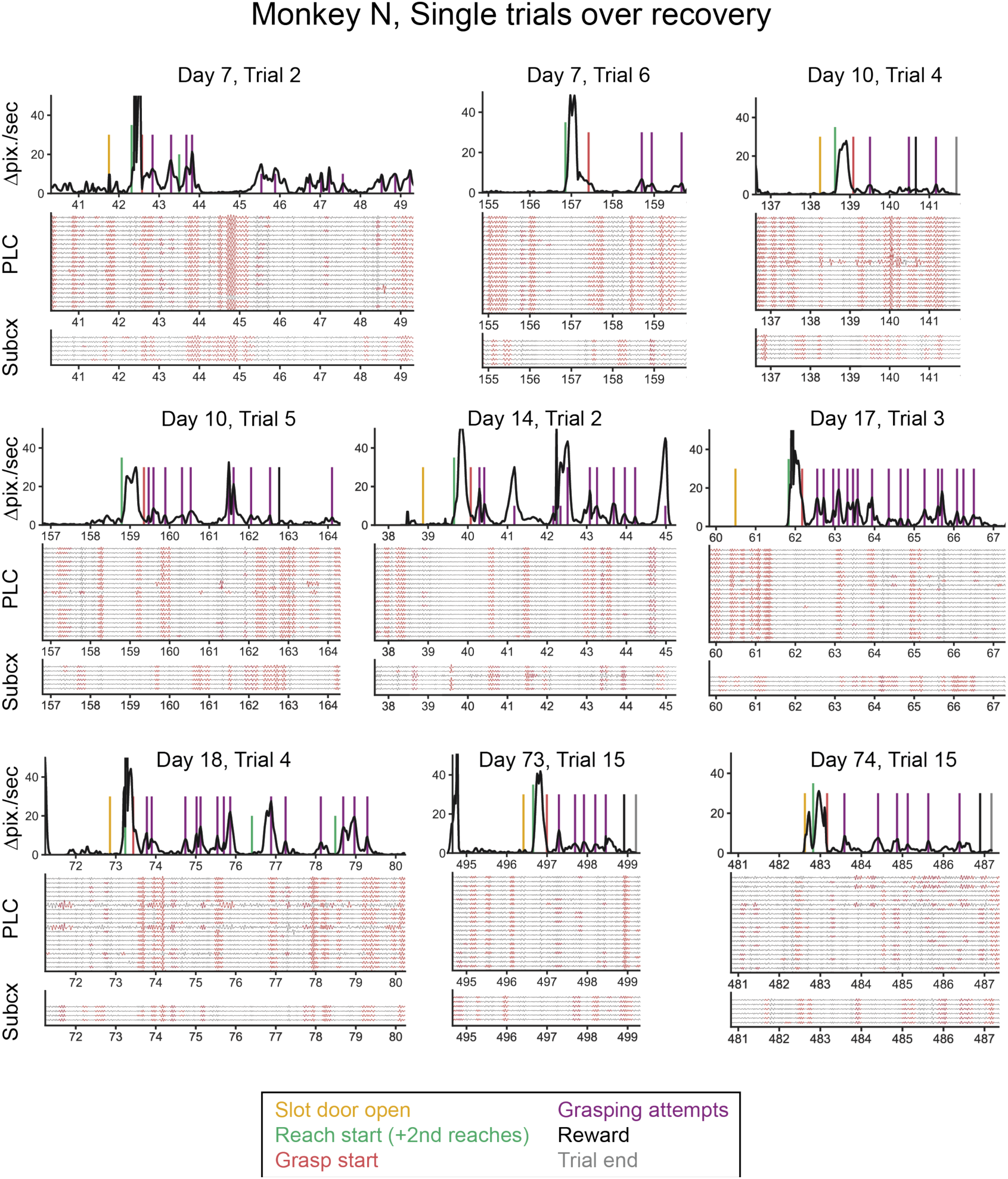
Monkey N single trials over the course of recovery. In the same style as Fig. 1JK.

**Figure S6.**
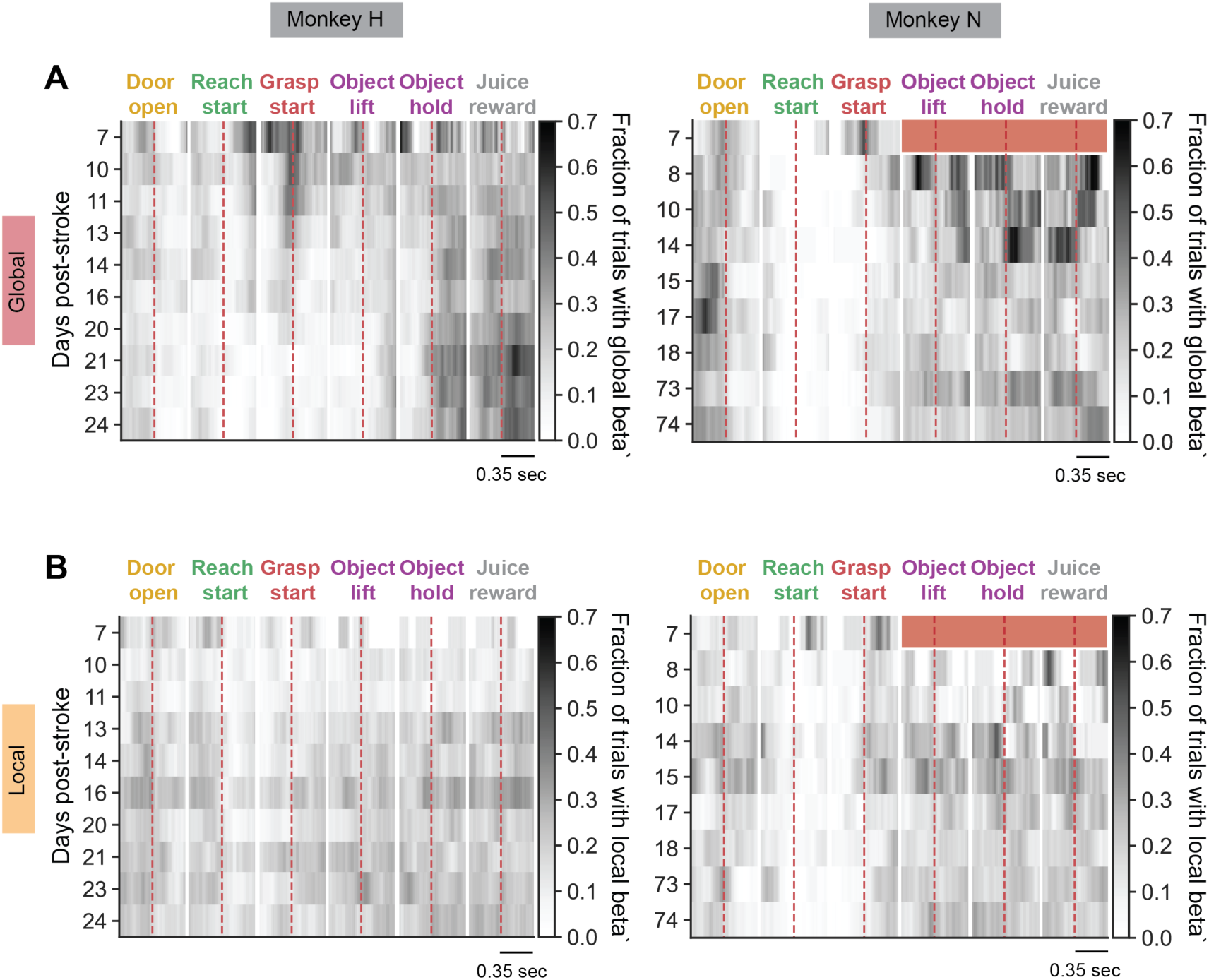
Alternative visualization of data in Fig 4AB. Individual post-stroke days are shown as rows, and darkness of the row indicates higher fraction of trials with global (A) or local (B). Red shading indicates no data present (no trials with successful lifts in Monkey N on day 7).

**Figure S7.**
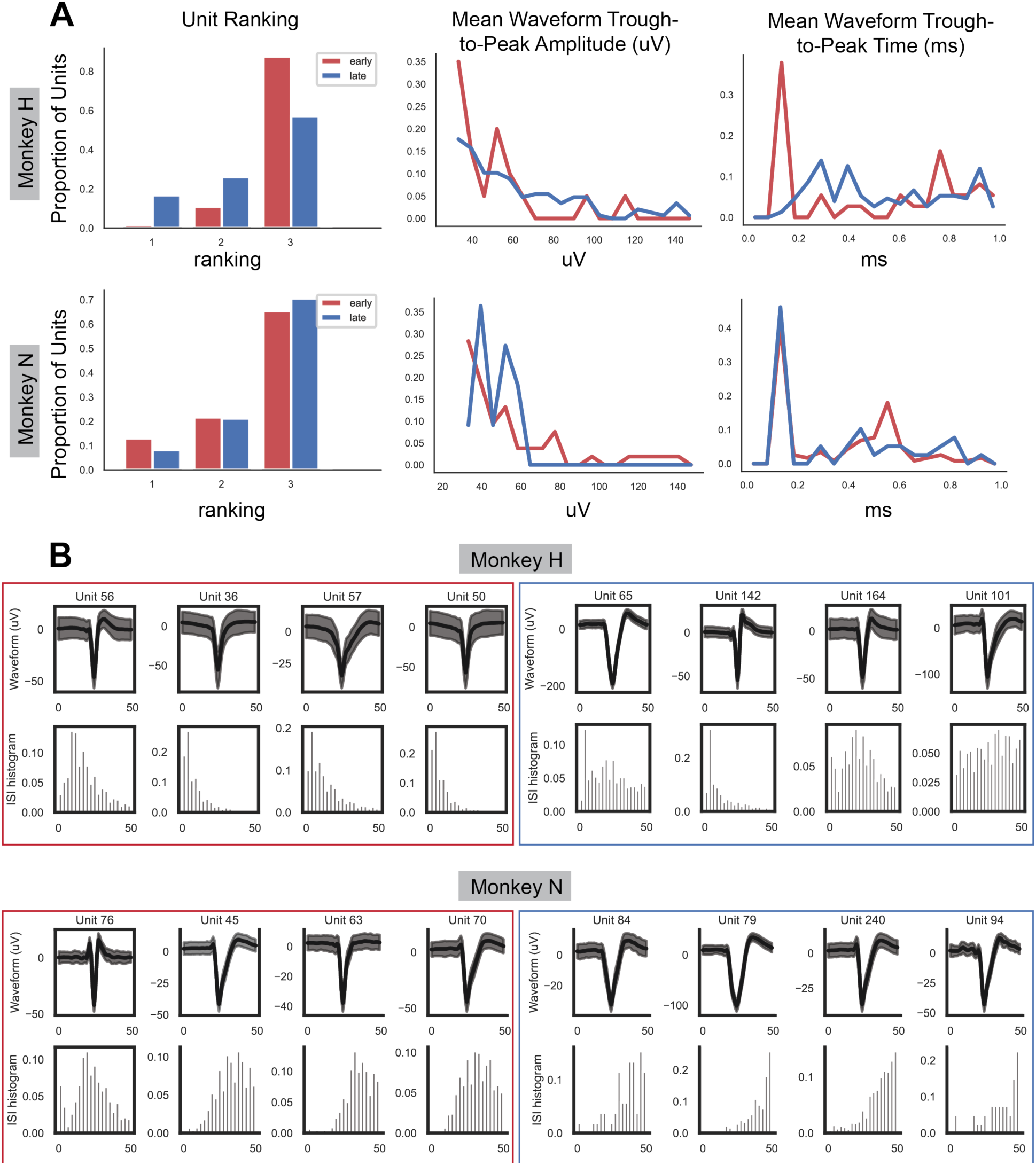
Properties of single units recorded during early recovery and late recovery. (A, *left*) Unit ratings during early vs. late recovery. Units were rated according to SNR: Rating 3: SNR > 3.0, Rating 2: SNR > 4.0, Rating 1: SNR > 8.0. Monkey N had very similarly ranked units early vs. late in recovery whereas Monkey H had a higher proportion of higher ranked units late in recovery. (A, *center*) Distribution of trough-to-peak amplitude (uV) for early vs. late units. Animals show trends in different directions: Monkey H has higher amplitude units later in recovery whereas Monkey N has higher amplitude units early in recovery. (A, *right*) Distribution of waveform widths (measured via trough-to-peak time) for animals. Monkey N has a very consistent distribution of waveform widths in early vs. late whereas Monkey H has a higher proportion of very short waveform widths early in recovery compared to late. (B). Examples of waveforms and inter-spike interval histograms from randomly selected units from Monkey H and Monkey N during early (red box) and late (blue box) recovery.

